# Cellular life from the three domains and viruses are transcriptionally active in a hypersaline desert community

**DOI:** 10.1101/839134

**Authors:** Gherman Uritskiy, Michael J. Tisza, Diego R Gelsinger, Adam Munn, James Taylor, Jocelyne DiRuggiero

**Author notes:** Corresponding authors and; Correspondence: Jocelyne DiRuggiero Johns Hopkins University Department of Biology, 3400 N. Charles Street, Mudd Hall 235 Baltimore MD 21218, USA.

## Abstract

Microbial communities play essential roles in the biosphere and understanding the mechanisms underlying their functional adaptations to environmental conditions is critical for predicting their behavior. This aspect of microbiome function has not been well characterized in natural high-salt environments. To address this knowledge gap, and to build a general framework of relating the genomic and transcriptomic components in a microbiome, we performed a meta-omic survey of extremophile communities inhabiting halite (salt) nodules in the Atacama Desert. We found that the major phyla of this halophilic community have very different levels of total transcriptional activity and that different metabolic pathways were activated in their transcriptomes. We report that a novel *Dolichomastix* alga – the only eukaryote found in this system – was by far the most active community member. It produced the vast majority of the community’s photosynthetic transcripts despite being outnumbered by members of the *Cyanobacteria*. The divergence in the transcriptional landscapes of these segregated communities, compared to the relatively stable metagenomic functional potential, suggests that microbiomes in each salt nodule undergo unique transcriptional adjustments to adapt to local conditions. We also report the characterization of several previously unknown halophilic viruses, many of which exhibit transcriptional activity indicative of host infection.

**Originality-Significance Statement:** While the metagenomics of hypersaline environments have already led to many discoveries, the transcriptional adaptations and functions of halophilic microbial communities in natural environments remains understudied. We perform the first robust meta-omic investigation of a hypersaline desert ecosystem, linking the genomic and transcriptional elements of the community. Our analysis unexpectedly revealed that Eukaryotes may be the main primary producers in this extreme environment, despite halophilic Archaea and Bacteria dominating the biomass. We also expand on the existing known diversity of halophilic viruses and demonstrate abundance (copies per million reads) and metatranscriptomic activity (transcripts per million reads); putative hosts are shown on the right.

## INTRODUCTION

While transcriptional and metabolic activities of human-associated microbiomes are being actively explored, relatively little is known about these activities in environmental microbiomes, particularly in hypersaline environments (Ramos-Barbero et al., 2018). Characterizing the metatranscriptome of a natural microbial community puts into perspective the relationship between its transcriptional function and functional potential based on metagenomic approaches, and give a more accurate view of the functioning of the community as a whole. For example, previous study of atmospheric microbiomes, comparative multi-omics showed that different taxonomic groups can constitute different fractions of metagenomic and metatranscriptomic components of the microbiome (Amato et al., 2019). A study of microbiomes in hot alkaline sulfur springs revealed that both aerobic and anaerobic organisms were simultaneously active in the springs and involved in sulfur and nitrogen cycling, suggesting microscopic sub-compartmentalization of obligate anaerobes within the community (Tripathy, Padhi, Mohanty, Samanta, & Maiti, 2016). In an alkaline lake, metatranscriptomic analysis of the communities inhabiting varying depths allowed identification of key pathways for to sulfur and arsenic cycling, and the organisms contributing to each process at varying depths (Edwardson & Hollibaugh, 2017). However, metatranscriptomics studies are difficult to conduct in most extreme environmental microbiomes due to low available biomass and high sample complexity (Ramos-Barbero et al., 2018).

Hypersaline microbiomes remain some of the most understudied from the transcriptomic perspective. To date, the study of hypersaline environments has greatly advanced our understanding of core microbiology principles, evolution, and the origin of life itself (Gunde-Cimerman, Plemenitas, & Oren, 2018; Oren, 2008; Paul & Mormile, 2017). Halophiles and their activities have been extensively studied for their unique adaptations, phylogenetic diversity, and potential economic and scientific benefits. Halophilic microbiomes provide opportunities for interdisciplinary activities and exploration of promising solutions for current and future global issues, particularly in the context of global warming and the search for life on other planets (Paul & Mormile, 2017). The metagenomic components of a number of high-salt natural microbiomes have been partially resolved, particularly from salterns and hypersaline lakes, where assembly and binning of shotgun sequencing data has led to the discovery of hundreds of novel taxa (Hedlund, Dodsworth, Murugapiran, Rinke, & Woyke, 2014; Ramos-Barbero et al., 2018; G. Uritskiy & DiRuggiero, 2019). However, the transcriptional and metabolic activities of such communities remain understudied (Ramos-Barbero et al., 2018).

Investigating the relationships between the metatranscriptomic and metagenomic components of partially segregated communities has the potential to offer unique insight into the deterministic and stochastic factors contributing to community function. While recent metagenomic studies, in isolated and taxonomically unique microbial communities, showed that such communities converge to similar functional potential landscapes (Louca et al., 2016; G. Uritskiy et al., 2019), it remains to be seen how this variance is reflected in the transcriptional landscape. Such studies have been conducted in human-associated microbiomes and it was found that functional potentials are only weakly correlated with the communities’ transcriptional functioning (Abu-Ali et al., 2018). In a hydrothermal vents multi-omic study, microbial communities were found to diverge at the transcriptional and the functional potential levels despite being subject to similar conditions and having a similar taxonomic structure (Fortunato, Larson, Butterfield, & Huber, 2018).

Extremophile endolithic (inside-rock) microbiomes of the Atacama Desert, Chile are a compelling ecology model for studying microbial community assembly principles and their sensitivity to perturbation make them an effective system to investigate the effects of climate change on microbiomes (G. Uritskiy et al., 2019). Encased in rocks, these communities have minimal exchange of biomass and nutrients with the outside, allowing each community to develop independently and, as such, providing insights into community structure replication. Despite an average annual precipitation of less than 1mm (Davila et al., 2015) microbes have evolved to survive by living inside halite nodules (salt rocks), relying on atmospheric moisture absorbed by the salt through deliquescence (Crits-Christoph et al., 2016; Robinson et al., 2015).

Based on functional potential, the only members of the halite community capable of carbon fixation are a single species of halophilic alga and 1-3 species of cyanobacteria (*Halothece* and *Euhalothece*). The carbon they fix also supports a number of heterotrophs, including *Halobacteria* and *Bacteroidetes*, which constitute the majority of the system’s biomass (Crits-Christoph et al., 2016; Robinson et al., 2015). The community was also found to contain a diverse consortium of viruses infecting most members of the Archaea and Bacteria domains (Crits-Christoph et al., 2016). Analysis of temporal (G. Uritskiy et al., 2019) and spatial (Finstad et al., 2017) dynamics of this community revealed water to be the governing factor of its taxonomic composition. It is also likely that the humidity and temperature changes in the Atacama Desert throughout the diurnal (day/night) cycle have an impact on microbiome functioning. A previous study demonstrated that these communities are metabolically active with *in situ* activity measurements for photosynthesis and respiration, however the contributing community members on the molecular level have not been identified (Davila et al., 2015). Our recent interrogation of the community’s small non-coding RNAs showed that these communities are capable of dynamic regulation of their transcriptome with sRNAs (Gelsinger et al., 2019).

To address the lack of knowledge on the transcriptional functioning of halophilic microbial communities, we interrogated the metatranscriptome of the halite microbiome. We sampled several nodules from different time-points, placing our transcriptional observations in the context of the metagenomic composition of the respective samples, and revealing the transcriptional activities in this unique community. We characterized the functional pathways transcriptionally active in the community, and found a surprising degree of transcriptional activity of the community’s only Eukaryote. We also reported on the phylogenetic variety and infective activity of this community’s viruses.

## RESULTS

### Community taxa varied in their contribution to the metatranscriptome

Genomic DNA and total RNA for 12 biological replicates, each from a different halite nodule, were sequenced, yielding a total of 96.0M and 81.2M reads, respectively. RNA reads matching rRNAs were removed computationally, yielding 12.8M non-ribosomal RNA reads. Assembled contigs were taxonomically classified based on genes annotated within then. The DNA and RNA reads were then aligned back to these classified contigs and the overall taxonomic composition at the metagenomic and metatranscriptomic level was determined from the distribution of the reads aligning to each respective taxonomic classification. (Fig. 1). This analysis revealed differences between the community taxa abundance (fraction of DNA reads) and overall gene expression levels (fraction of RNA reads), as the transcriptional level of several taxa were over-or under-represented with regard to their relative abundance in the community. In particular, the total transcriptional activity of Eukarya, represented by a single metagenome-assembled genome (MAG; see following sections), was much higher than their genomic abundance in the community; 12% of non-rRNA metatranscriptomic reads were mapped to Eukaryotic contigs, compared to only 1% of metagenomic reads. In contrast, bacteria – particularly *Bacteroidetes* – represented a smaller fraction of the metatranscriptome compared to the metagenome (8% vs 15%).

**Fig. 1:**
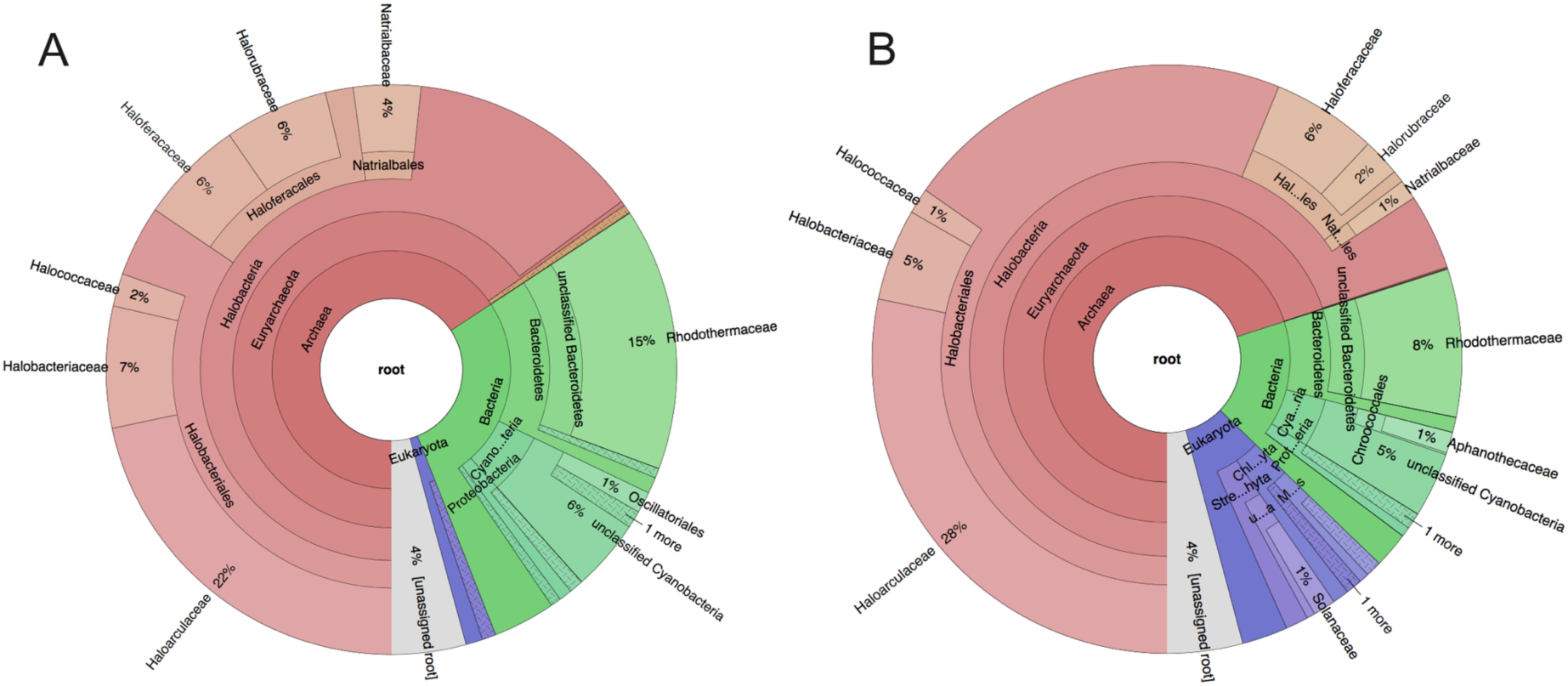
Taxonomic composition of the halite community estimated from the metagenome (A) and metatranscriptome (B). Relative compositions were calculated from the read coverage of contigs annotated to each taxon. The contigs were first classified based on the phylogenetic lineages of the genes that they carried, estimated with the IMG annotation service.

To understand the relationship between relative abundance of organisms and their transcriptional activity, we carried out an analysis at the level of individual contigs. The relative abundance of contigs, expressed in copies per million (CPM), was estimated from their DNA read coverage and the contig transcriptional activity was estimated from the average expression of genes carried on the respective contigs, expressed in transcripts-per-million (TPM). We found that contigs from different taxa displayed drastically different transcriptional activity levels, which were not necessarily correlated with their abundance levels (Fig. 2A). In particular, the three photosynthetic species – *Halothece* and *Euhalothece* cyanobacteria and a *Chlorophyta* green alga – had similar average gene transcriptional levels despite being present at notably different genomic abundances. Contigs from the dominant cyanobacteria, *Halothece*, had high gene expression levels but, because of the high relative abundance of this organism in the community, the transcriptional activity of the contigs relative to their abundance was low. As previously observed in the overall taxonomic composition of the community, the algae contigs displayed extremely high levels of transcriptional activity despite being present at very low genomic abundance (∼1%). On the other extreme, *Nanohaloarchaea* had very low gene transcription levels and low relative abundance. The differences in abundance and transcriptional activity levels of other taxa were not as extreme, however significant variation in transcriptional activity was observed within the highly diverse groups of *Bacteroidetes* and *Halobacteria* (the dominant class within the *Euryarchaeota* phylum).

**Fig. 2:**
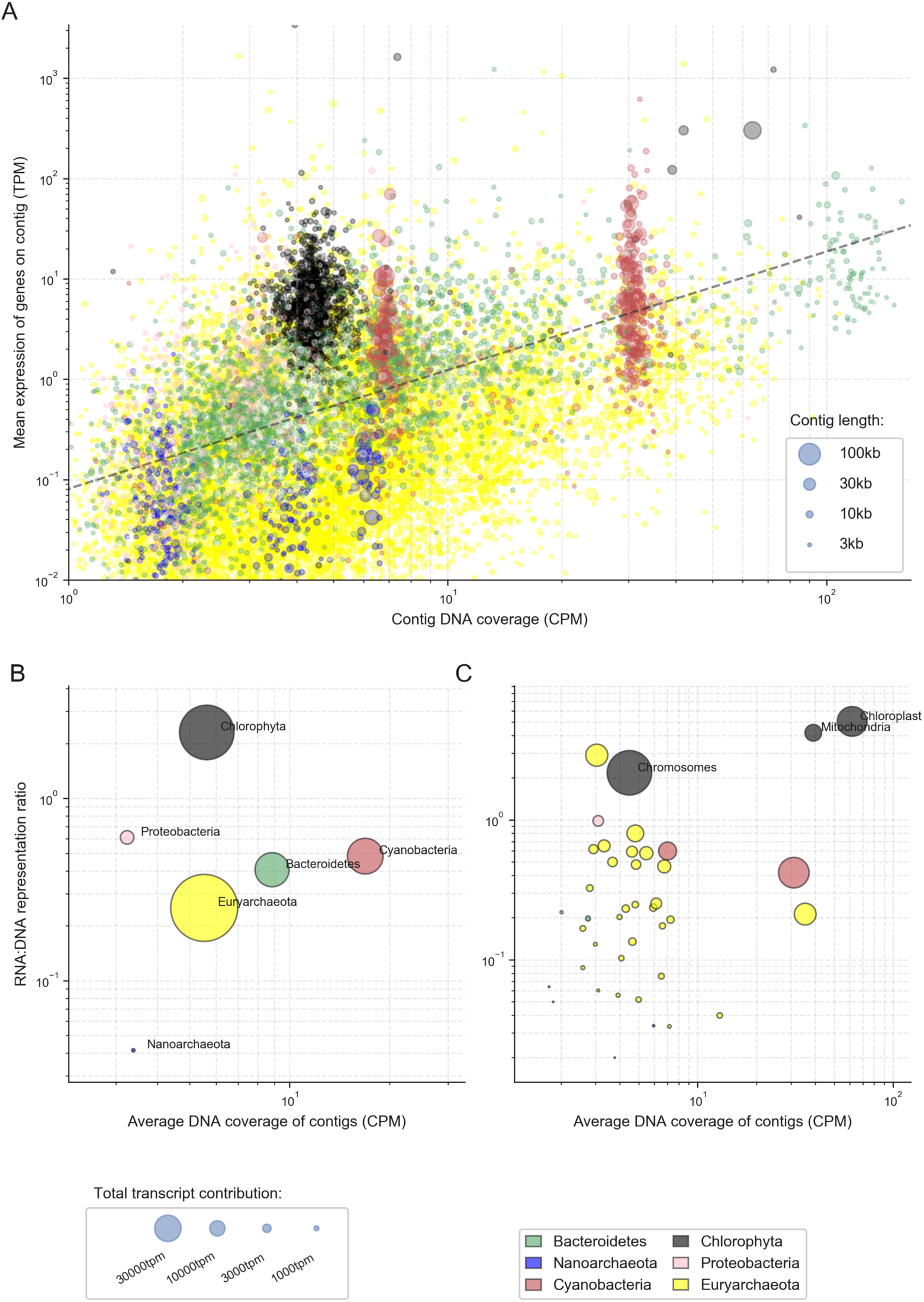
Transcriptional activity vs abundance of contigs and MAGs. (A) Total transcriptional activity (y-axis) of contigs in relation to their abundance in the metagenome (x-axis). Relative transcriptional activity (y-axis) of contigs (B) and MAGs (C) shown in relation to their average relative abundance (x-axis) and total contribution to the transcriptome (circle size). The total transcriptional activity of a given contig or MAG was estimated to be the median read expression (TPM) of its genes. The total relative abundance was estimated from its average DNA read coverage.

Quantifying the relationship between relative abundance and transcription levels at the phylum level (Fig. 2B) or using metagenome-assembled genome (42 MAGs, completion>70%, contamination<5%; Data 1; Fig. 2C) supported that the *Chlorophyta* (algae) was an order of magnitude more active than most other MAGs in this community. In terms of the total number of transcripts (Fig. 2B, 2C), algae contributed more to the metatranscriptome than both of the other photosynthetic members combined. While alga’s organelle genomes were much more numerous relative to the chromosomal genome (approximately 10X for mitochondria and 20X for chloroplasts), we found that, even when adjusting for genome copy numbers, the chloroplasts were nearly 10X more transcriptionally active than the *Cyanobacteria* (Fig. 2C). The alga was taxonomically identified to be in the class *Mamiellophyceae*, whose members have been previously characterized to have a single mitochondria and chloroplast (Robinson et al., 2015; van Baren et al., 2016). This suggests that the coverage differences between the halite alga’s main chromosomes, mitochondria, and chloroplasts were the result of multiple genome copies in each organelle.

While some taxa were more transcriptionally active than others, we also found that the transcriptional activity of MAGs (expressed as the ratio of total RNA expression to the total DNA abundance) changed significantly between replicates (Fig. S2). This variance in activity was so great that an attempt to correlate the abundance and transcriptional levels of any given MAG across replicates (Fig. S3C) was successful for only a small subset of organisms (Fig. S3D). We verified that these findings were not the results of binning biases by repeating the analysis at the contig level (Fig. S3A, B).

### Virus diversity and transcriptional activity

We investigated viral diversity and transcriptional differences in the halite community using the sequences of homologous viral proteins, across samples. We extracted and functionally annotated 91 viral contigs from the metagenome, ranging in size from 5Kbp to 51Kbp. The halite viruses included members of order *Caudovirales* (prokaryotic head-tail viruses including families *Myoviridae*, *Siphoviridae*, *Podoviridae* and so-called *Haloviruses*) and of the archaeal virus families *Pleolipoviridae* and *Sphaerolipoviridae* (Fig. S4). The extracted viral contigs represented expansions of some extant genera (as determined by VContact2) while a number of contigs established previously unidentified genus-level clades. CRISPR spacer and nucleotide sequence similarity between viral genomes and host chromosomal regions were used to identify putative hosts of prokaryotic viruses, leading to the identification of the hosts for 44 viruses, and revealing that the viruses primarily targeted *Halobacteria* and *Salinibacter* hosts for infection. The most abundant and transcriptionally active viruses belonged to a new clade of *Myoviridae* viruses, predicted to infect members of the *Halobacteria* class (Fig. 3). Interestingly, most highly transcribing viruses were active in only a few samples. The most highly expressed genes were annotated as being virion structural components or DNA replication/mobilization, indicating that active lytic infection was occurring in these communities at the time of sampling.

**Fig. 3:**
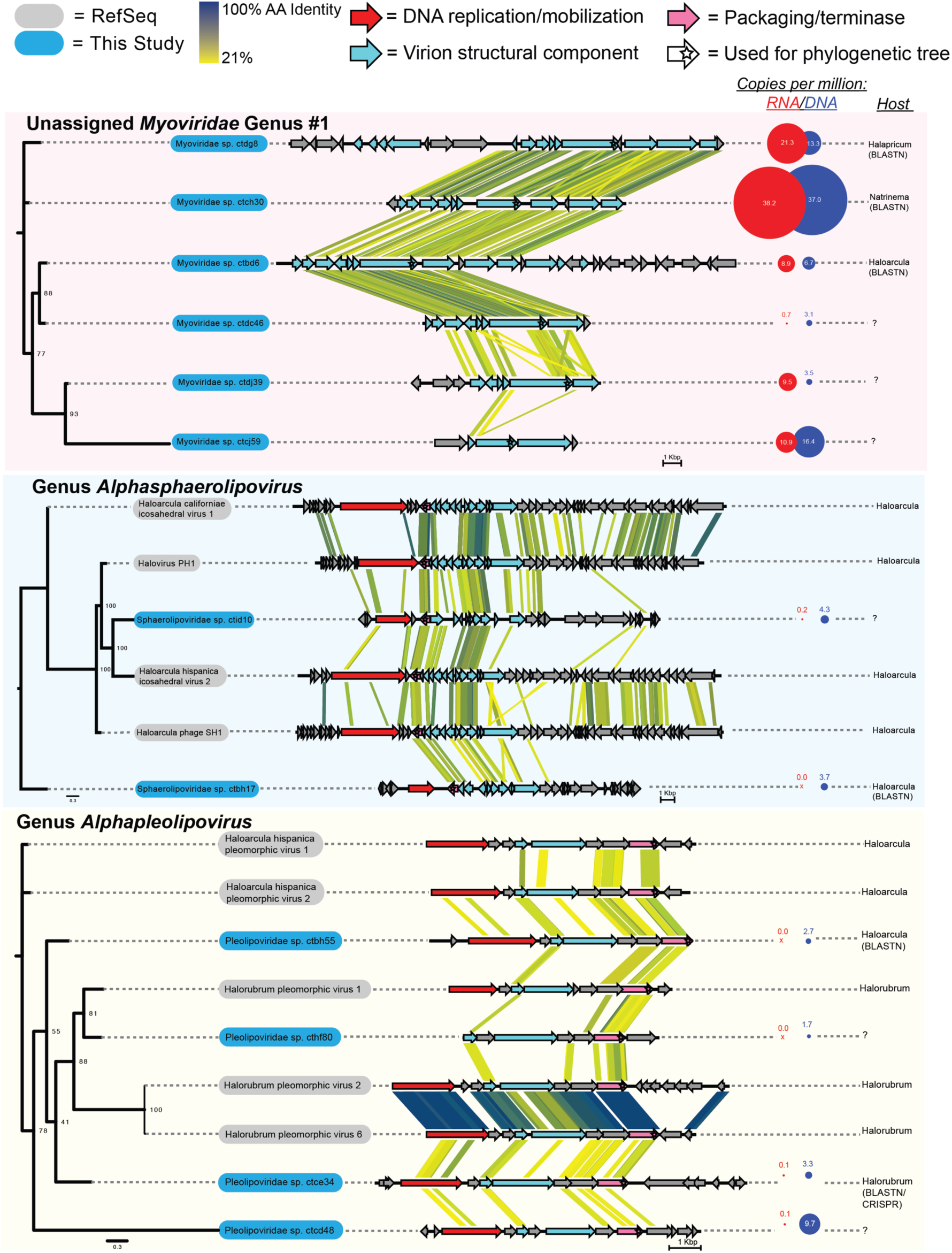
Diversity of viruses in the halite community. Viral contigs from this study (in blue) annotated to a novel Myoviridae genus cluster and to Alphaspaerolipovirus and Alphapleolipovirus genera, and viral contig from the RefSeq database (in grey). Genome maps were drawn and compared with TBLASTX and EasyFig. Circles represent metagenomic

In order for a virus to be transcriptionally active it needs intracellular access to a host cell, suggesting that the activity level of a given virus in a sample is dependent on its abundance and transcription rates, but also on the abundance of its host and its infection success. Consistent with this, we found that a virus’s high genomic abundance does not necessarily lead to transcriptional activity, and most viruses had no detectable transcription in most samples (Fig. S5). In a given sample, many transcriptionally active viral contigs were not highly abundant at the DNA-level, and highly-abundant contigs were not necessarily active at the transcriptional level. Only 12 out of the 91 virus contigs showed significant positive correlation between viral abundance and viral transcriptional activity (Fig. S6).

### Characterization of putative genome of a novel *Dolichomasti*x alga

Through a combination of assembly and binning methods we reconstructed the alga chromosome, chloroplast, and mitochondrion draft genomes. The metatranscriptome was also assembled to obtain the complete sequence of the 18S rRNA gene sequence. The latter sequence was found to be 99% identical to the sequence previously extracted from the halite microbiome using an amplicon-based approach (Robinson et al., 2015), and 94% identical to a putative *Dolichomastix* alga found in Lake Tyrell (Heidelberg et al., 2013). Re-evaluation of the phylogenetic placement of this sequence confirmed that the halite alga belongs to the *Dolichomastix* genus (Fig. S7).

Evaluating the relative contributions of oxygenic photosynthetic members to the functioning of the community revealed that the halite alga contributed significantly to photosynthesis. We identified several key genes from the photosynthetic pathway, a set of which were found in all three organisms capable of photosynthesis, the alga *Dolichomastix* and the two *Cyanobacteria* (Fig. 4). For almost every photosynthetic gene – particularly *psa* genes and most *psb* genes – the algae contributed an order of magnitude more transcripts than both of the *Cyanobacteria* combined. We also identified components of the carbon fixation pathway in all three phototrophs and found that the chloroplast gene encoding for RuBisCO was the second-highest expressed non-ribosomal gene in the entire alga (after *psb*D).

**Fig. 4:**
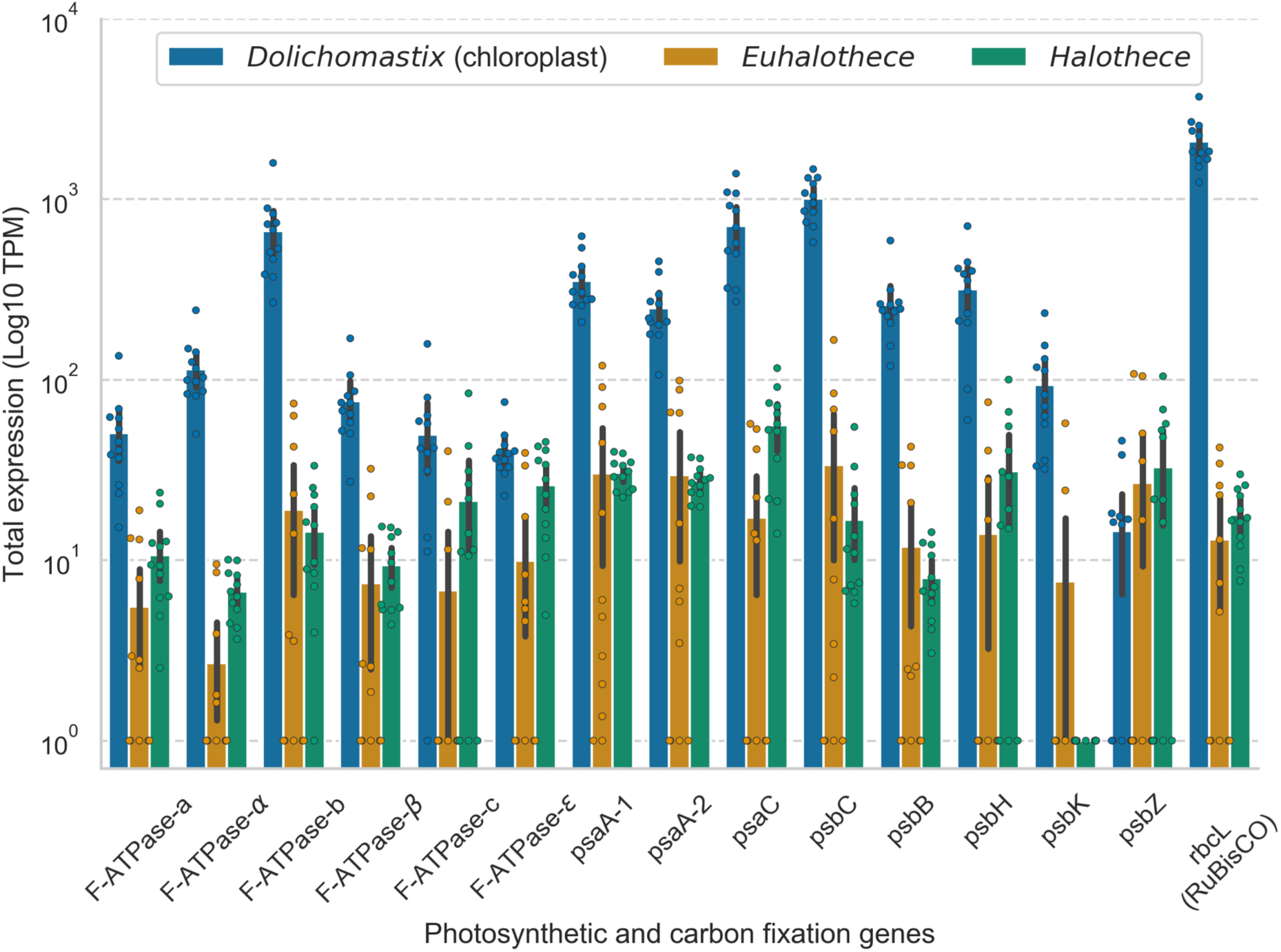
Expression of photosynthetic and carbon fixation genes identified in the genomes of the 3 organisms capable of oxygenic photosynthesis in the halite community.

Given the high importance of *Dolichomastix* alga for community carbon fixation, we further characterized its genome in relation to its closest fully sequenced phylogenetic relatives *Ostreococcus tauri* and *Micromonas pusilla* (marine algae), and another halophilic alga *Dunaliella salina*. The extracted *Dolichomastix* chromosomal genome was 10.3Mbp long, compared to 13.1Mbp in *O. tauri*, 22.4Mbp in *M. pusilla*, and 343.7Mbp in *D. salina*. The chloroplast and mitochondrion genomes of the halite alga were 95Kbp and 46Kbp, respectively, but non-chromosomal sequences of closely-related algae were unavailable for comparison. Using BUSCO (Seppey, Manni, & Zdobnov, 2019), the *Dolichomastix* genome was evaluated to be 45.3% complete with 0.2% contamination, however given the lack of highly compact eukaryotic genomes in the BUSCO database, the completion value might be underestimated. All four genomes were functionally annotated with the same method and the isoelectric point (p*I*) for each predicted protein was estimated (Fig. S8A). We found that the pI distribution for the predicted proteome of the *Dolichomastix* alga was significantly different from that of *D. salina*, but relatively similar to its non-halophilic closest phylogenetic relatives *M. pusilla* and *O. tauri*. However, the non-halophiles displayed a trimodal pI distribution curve, with a notable fraction of the genes predicted to encode for proteins with a pI of 10.5 or greater. *Dolichomastix* on the other hand, lacked this third peak, similar to the more distant but halophilic alga *D. salina*. Of the few *Dolichomastix* proteins with a high predicted p*I* (p*I*>11) were histone and DNA-associated proteins, which need to be alkaline to function.

To investigate whether the genes encoding for high pI proteins in *M. pusilla* and *O. tauri* were present in the halite *Dolichomastix*, we aligned the amino acid sequences of the *M. pusilla* and *Dolichomastix* genomes. Of the 105 *M. pusilla* genes encoding for high-p*I* proteins that were found to have homologues in *Dolichomastix*, 80 encoded for proteins with a significantly lower p*I* in *Dolichomastix* (Fig. S8B). These proteins were not significantly enriched for any specific pathway or cellular compartment, but many were chaperones, mRNA processing proteins, and DNA-binding proteins.

To further compare the halotolerant *D. salina* and *Dolichomastix* genomes, we investigated homologs of important high-salt adaptation proteins. Previous proteomic studies of *D. salina* identified 51 cellular and 46 membrane-bound proteins that were upregulated under high-salt stress (Katz, Waridel, Shevchenko, & Pick, 2007; Liska, Shevchenko, Pick, & Katz, 2004). We mapped these genes to the *Dolichomastix* genome and found 33 homologous genes with a potential role in high-salt tolerance. Of these, all were actively expressed in *Dolichomastix* and a significant majority were highly expressed (>10TPM, Fig. S8C). Using this homologous set of genes with potential roles in high-salt tolerance in *D. salina*, we identified several highly expressed chaperones and heat-shock proteins in the *Dolichomastix* transcriptome, as well as a number of mitochondrial and chloroplast genes (Data S2). The predicted proteome in the chloroplast and mitochondria generally favored high-*pI* proteins compared to the main cellular compartment, although the small number of organelle genes may bias the observed *pI* distributions, particularly in the mitochondria (Fig. S8D).

### Major functional pathways highly expressed in the community

Investigating the expression levels of functional pathways from the KEGG Brite database in the halite community allowed us to identify highly transcribed functions. The transcriptional activity (transcripts per million reads, or TPM) of pathways was estimated from the sum of expression values of all the genes in a pathway, and standardized to an equal sum of pathway TPMs in each replicate. Interestingly, the Euclidian distance hierarchical clustering between RNA replicates based on pathway expression did not reflect the sampling time-points or the taxonomic compositions of the samples (Fig. 5). Most highly-expressed pathways, including translation, nucleotide metabolism, amino acid metabolism, and DNA replication and repair, were also present at high levels in the functional potential (copies per million of DNA reads, or CPM). In contrast, photosynthesis was the most highly expressed pathways in the community but had relatively low levels of abundance at the DNA level, which were similarly estimated from the total DNA read coverage of genes in each pathway (see methods). Opsin production pathway was also expressed at relatively high levels when taking into consideration its low abundance in the functional potential.

**Fig. 5:**
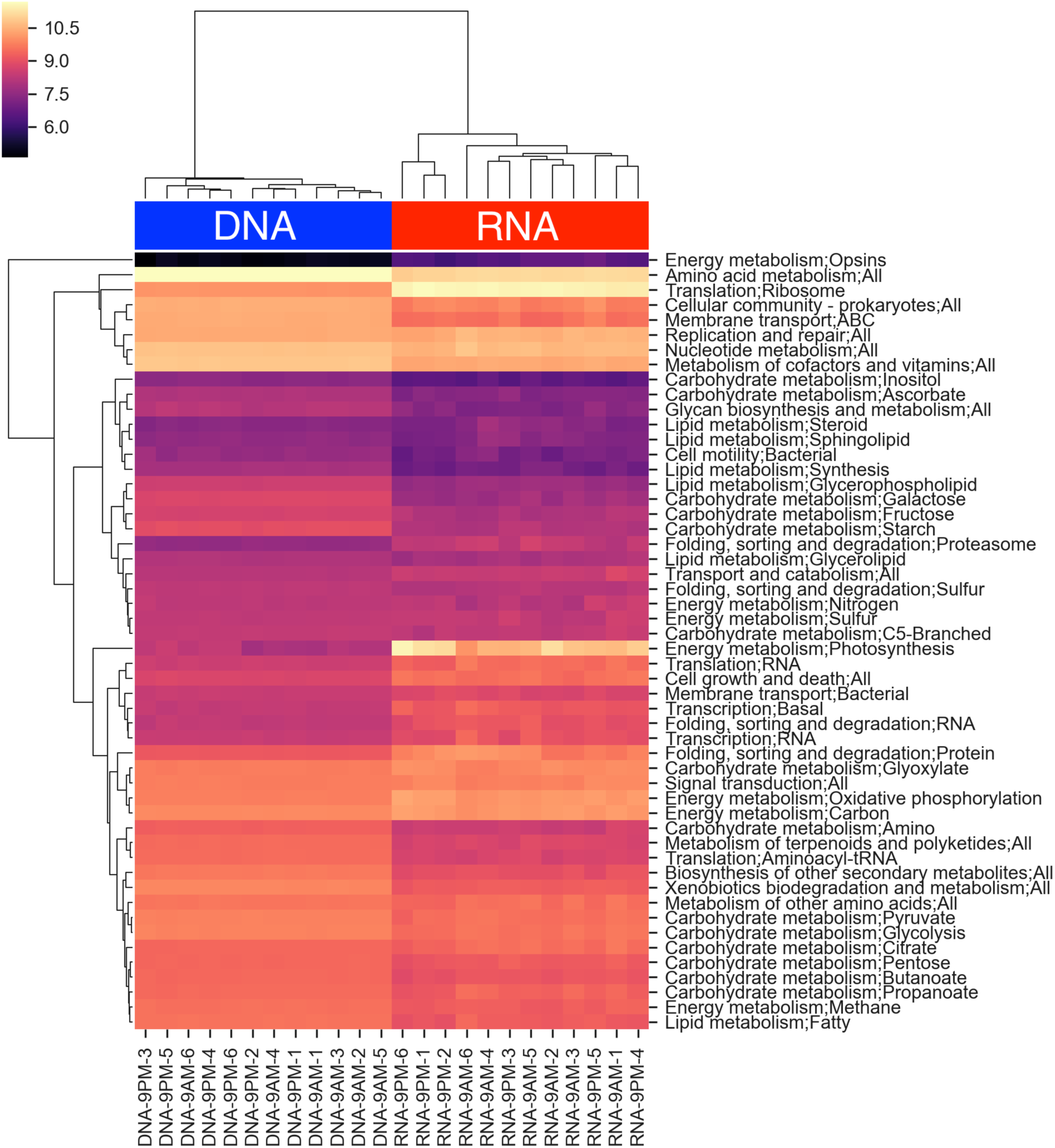
Relative abundance and expression of major KEGG functional pathways in the metagenomic and metatranscriptomic samples. The pathway abundance (DNA) and expression (RNA) value were the combined total CPM or TPM (respectively) of the genes that constitute that pathway. All samples were standardized to an equal total coverage of pathways.

By standardizing the abundance and expression values of each pathway to the maximum value (abundance or expression), we were able to visually compare the relative expression of each pathway in relation to its abundance in the metagenome and infer pathways transcriptionally prioritized by the community (Fig. S9). The Euclidian hierarchical clustering of community metabolic pathways resulted in three groups – those that were expressed lower, higher, or equal to their respective abundances in the community’s functional potential. Among the lowly-active pathways, we found cellular membrane and cell wall components, including most lipid metabolism and synthesis pathways, and glycan biosynthesis. Within the highly expressed pathway group we found functions responsible for energy metabolism and carbon flow, such as photosynthesis and oxidative phosphorylation. The other group of highly expressed pathways dealt with protein turnover. In particular, multiple pathways for transcription and translation were very active, as were pathways for protein folding, trafficking, and degradation.

### Pathway expression enrichment in major taxa

We further explored the pathways most transcribed for each of the major taxonomic groups in the community, *Halobacteria, Bacteroidetes, Chlorophyta* (green algae), *Cyanobacteria*, *Proteobacteria, Actinobacteria, and Nanohaloarchaea*, by computing the ratio between the standardized pathway expression to pathway abundance (in the metatranscriptome and metagenome, respectively; Fig. S10). Importantly, these ratios did not represent a taxon’s overall functional landscape, but rather the degree to which a specific pathway was expressed.

While each major taxon in the community had a unique transcriptional profile, we found many similarities between their highly expressed pathways. For all major taxa, except *Nanohaloarchaea*, protein folding, sorting, and degradation pathways were the most highly expressed pathways. General carbon metabolism and oxidative phosphorylation were also highly expressed, although the specific sugar metabolism pathways were highly varied. The community’s least active members – the *Nanohaloarchaea* – where the most different from the other community phyla. Their only highly-expressed pathways were for nucleotide metabolism, transcription and translation as well as pathways for metabolizing simple organism molecules such as fructose and pyruvate. Alignments with BLAST were used to identify fragments of the SPEARE gene encoding a protein essential in *Nanohaloarchaea* docking to their hosts (Hamm et al., 2019) in two of the MAGs (T17_Nanohaloarchaea_45_3 and T17_Nanohaloarchaea_46_6; Data S1), however none of these genes were expressed in the metatranscriptome.

Not surprisingly, the two oxygenic photosynthetic groups in the community – *Chlorophyta* and *Cyanobacteria* – clustered together in their transcriptional activity. They were the only taxa to carry and express the photosynthesis pathway, which was highly active in both. However, expression profile for other metabolic activities were quite distinct, with *Chlorophyta* strongly prioritizing glyoxylate metabolism and *Cyanobacteria* prioritized fructose and pentose metabolism. The opsin biosynthesis pathway, which produces the light-driven proton pumps in *Halobacteria* (bacteriorhodopsin) and *Bacteroidetes* (xanthorhodopsin), was one of the most highly expressed pathways in the metatranscriptome compared to the metagenome and was predominantly expressed in *Halobacteria*.

### No transcriptional differences detected between daytime and nighttime

Samples for metatranscriptomic (and the corresponding metagenomic samples) were collected at 2 time points during the diurnal cycle, at 9 am and 9 pm with 6 replicates, each from a different halite nodule, to uncover temporal transcriptional adaptations of community members. The 9 am time point was characterized by bright light (1h after sunrise) and high relative humidity (60%-80% RH), while the 9 pm time point was collected in the dark (3 hours after sunset) and at low RH (40%-50%). Differential expression analysis was performed on the entire community, standardizing the gene expression to the abundance of its contig in the DNA or the total contig expression in the RNA (Fig. S11A). DESeq identified differentially expressed genes, however the false discovery rate was greater than 5%, and the over-and under-expressed genes belonged to a seemingly random set of pathways. This analysis was also repeated for individual high-quality MAGs (>70% completion, <5% contamination), including the *Cyanobacteria* and *Dolichomastix* MAGs (Fig. S11B,C). Doing so allowed for a more robust standardization scheme that accounted for the abundance (or total expression) of the entire organism, but did not yield any significant differentially expressed genes. We were also unable to detect significant differences in total pathway expression (Fig. 5).

### The functional profile was more variable than the functional potential

To address the high inter-replicate metatranscriptome variation we observed in this work and investigate how pathway abundance and pathway transcription differences contribute to this heterogeneity, we computed the variance of KEGG pathway abundance or expression values across replicates, standardized to range from 0 to 1. This transformation allowed for direct comparison of the variance in pathway abundance to the variance in pathway expression in the whole community or for a taxon of interest. Across all of the community pathways we found a greater variation in pathway expression levels than pathway abundances (Fig. 6). This was true for all tested major taxa in the community (Fig. S12), suggesting that while the overall functional potential of the community remained relatively stable between replicates, consistent with previous findings, their transcriptional activity changed substantially in individual halite nodules.

**Fig. 6:**
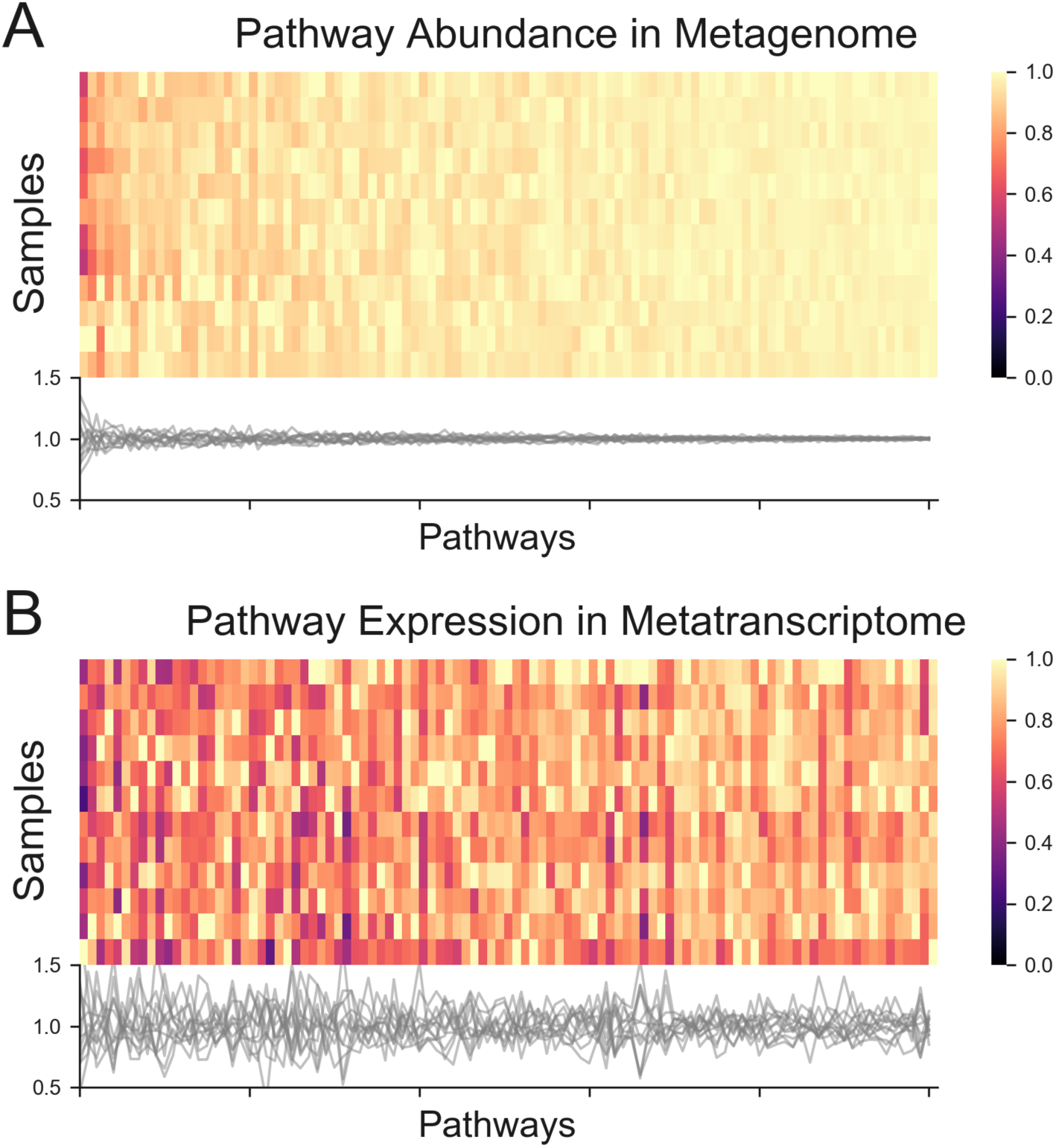
Inter-replicate variance in the metagenomic functional potential and metatranscriptome. KEGG pathway (A) CPM abundance and (B) TPM expression in sample replicates, standardized to the highest value in each column. Tracks below the heat maps show the CPM of TPM values from individual replicates, standardized to the average between all the replicates.

## DISCUSSION

Our analysis of both the metatranscriptomic and metagenomic components of a halite microbial community provided new insights into the functioning of this unique hyper-saline ecosystem. While previous research revealed some variation in inter-nodule taxonomic composition and functional potential (G. Uritskiy et al., 2019), here we found even greater inter-nodule differences in the metatranscriptomes. The heterogeneity we observed in metatranscriptomes most likely stems from a combination of varying rock topology, humidity and temperature metrics, which we could not measure at the time of sampling due to practical limitations. Previously, water availability has been described as the major factor driving community assembly in such arid environments (Finstad et al., 2017; Meslier et al., 2018), and it is likely that internal relative humidity at a given point in time varies significantly from nodule to nodule depending on their topology, exposure to sun and wind, and on their individual capacity for water retention. It has been shown that halite endoliths obtain most of their liquid water via salt deliquescence and that halite nodules progress through wet and dry stages during diurnal cycles (Davila et al., 2008). At air RH above 75%, mostly during the night and early morning, brine is formed inside the nodule, filling the pore space between halite crystals. During the day, water slowly evaporates from increasing temperatures and dry winds, leaving most of the nodule to dry. While the water dynamics inside each nodule is unique to that nodule, the overall conditions, and therefore the selective pressure from the environment, are very similar across all nodules, explaining the relative convergence of functional potentials of the halite communities’ pan-genomes.

We know from previous studies using pulse-amplitude modulation (PAM) fluorometry and respiration measurements that the halite communities are metabolically active, with significant differences between day and night (Davila et al., 2015). Transcriptional studies of aquatic cyanobacteria showed distinct temporal regulation of the photosynthesis and carbon fixation pathways in response to light (Saha et al., 2016; Welkie et al., 2019), with light-harvesting photosynthetic genes being upregulated early in the light period, and the core oxidative pentose pathway being activated during the dark period. We were unable to detect such changes in the transcriptome of the halite community, most likely because inter-halite transcriptional variations and a low signal to noise ratio.

Inter-cellular interactions, viral infection, and other stochastic biotic factors also likely influence the transcriptional landscapes found in each halite at any given point in time, compounding the complexity of linking the metatranscriptomic adaptations to specific factors. These results resemble the conclusions of an individualized gut microbiome multi-omic screening, in which the functional potentials and metatranscriptional landscapes of gut communities were only loosely correlated (Abu-Ali et al., 2018). These differences were detected both in the overall functional landscape of the community as well as in the transcriptional activities of individual genomes.

Comparing the taxonomic composition of the metagenomic and metatranscriptomic elements of the halite community revealed that many organisms have a drastically different contribution to the community’s functions than what could be inferred from previously reported metagenomic composition (Crits-Christoph et al., 2016). Past studies placed emphasis on cyanobacteria as major players in this microbial community, however we report that they are not the most transcriptionally active. Such results have also been found in other microbial communities with characterized metatranscriptomic components, including extremophile (Fortunato et al., 2018) and human-associated microbiomes (Franzosa et al., 2014), where many dominant taxonomic groups in the metagenomes were not the more transcriptionally active taxa. One of the least transcriptionally active members in the halite microbiome – *Nanohaloarchaea* – had minimal and sometimes undetected levels of transcription. Recent characterization of *Nanohaloarchaea antarcticus* in Antarctica suggests that they are ecto-parasitic, using a SPEARE protein complex to dock and form a feeding pore with a host *Halorubrum lacusprofundi* cell (Hamm et al., 2019). The low overall transcriptional levels of the halite *Nanohaloarchaea*, as well as undetectable expression of the SPEARE proteins, are indicative of their inactive state, most likely because they were not associated with a host at the time of sampling.

The most transcriptionally active organism in this community was an alga – the only eukaryote. Previous studies have identified eukaryotic components in hypersaline environments, notably green algae and protozoans (Harding & Simpson, 2018; Heidelberg et al., 2013; Robinson et al., 2015), however none reported their transcriptional or metabolic contribution to community functioning. The novel halophilic *Dolichomastix* alga was responsible for producing nearly 12% of the community’s non-ribosomal transcripts despite representing only 1% of the community’s metagenome and it was nearly 10 times more transcriptionally active than all the oxygenic photosynthetic prokaryotes. The relative transcriptional rates of the *Dolichomastix* alga and its organelles were still extremely high even when standardizing to genome size and gene count. The larger cell size of green algae compared to bacteria and archaea, and a higher basal metabolism required to survive in a high-salt environment, likely explain this novel finding. Broad estimates of the number of mRNA transcripts per average cell suggest 10^3^-10^4^ mRNA per bacterial cell and 10^5^-10^6^ mRNA for a 3000 µm^3^ eukaryotic cell (Ron Milo, 2016). Using previous microscopy-based observations of the halite alga, its volume can be roughly estimated to be roughly 1000 µm^3^, assuming a spherical cell shape (Robinson et al., 2015). Taken together, these estimates suggest that the halite alga is expected to be 10-100 times more transcriptionally active than the prokaryotic species, which is similar to what is observed in this study. This suggests that the transcriptional overrepresentation of the alga in this community likely stems from the cell size and functional differences between the eukaryotes and prokaryotes rather than activity levels. This representation disparity is similar to what was found in a cow rumen microbiome, where a multi-omic study revealed that a small eukaryotic minority in the microbiome produced a significantly greater fraction of transcripts than expected from genome copy numbers (Comtet-Marre et al., 2017). We also found that the alga is responsible for producing the vast majority of the photosynthetic transcripts in the system, which could suggest that they are the major primary producers of the community. In a previous study of coastal sediments exposed to light, eukaryotic diatoms were shown to dominate the community metatranscriptome, particularly with extremely high numbers of photosynthetic pathway transcripts (Broman, Sachpazidou, Dopson, & Hylander, 2017).

The novel *Dolichomastix* MAG showed transcriptional adaptations similar to a model halotolerant algae *D. salina*. Previous studies with *D. salina* reported the upregulation under high salt of major metabolic pathways involved in photosynthesis and carbon fixation to produce enough energy and glycerol-based secondary metabolites to actively balance high external salt concentrations (H. Chen & Jiang, 2009). Similarly, we found that *Dolichomastix* had extremely high rates of photosynthesis and oxidative phosphorylation, likely producing secondary metabolites to counter-act external osmotic pressure (Oren, 2014; Polle et al., 2017). However, we were unable to detect evidence for active glycerol production in the alga’s transcriptome. The predicted proteome for *Dolichomastix* exhibited a lack of a high-p*I* proteins present in its non-halophilic phylogenetic relatives. While p*I* distribution in eukaryotes is not indicative of function, it has been linked to cytoplasmic and nuclear pH (Elevi Bardavid & Oren, 2012), suggesting that the *Dolichomastix* alga might have a slightly different intracellular environment compared to non-halophilic members of its class.

Our characterization of the viruses of the halite microbiome significantly expanded its existing sequence pool of viral diversity. Our shotgun meta-omic approach to virus and host discovery built on existing microarray-based research on halophilic virus metatranscriptomes (Santos et al., 2011), providing a basis for more homology-based discovery for viruses in other halophilic environments. Expanding on our previous metagenomic work (Crits-Christoph et al., 2016), we were able to reconstruct 3 times more viruses, many of which belonging to novel putative genera, including a wide variety of viruses targeting *Halobacteria* and *Salinibacter* hosts. We also detected significant transcriptional activity of genes encoding for viral structural and replicative components in the majority of the discovered viruses, indicating active infection of bacterial and archaeal hosts. Many viruses, including a *Halobacteria* virus from the *Myoviridae* class, displayed very high gene expression values, suggesting that they may play a significant role in shaping the structure and composition of halite communities. Our analysis of these viruses also revealed that their transcriptional activity (and thus active infection) did not correlate with their relative abundance nor with that of their host, indicating that their virulent success was dependent on a combination of deterministic (host and virus abundance) and stochastic factors (infectivity rates, host resistance accumulation) in any given halite nodule. These results are consistent with those of an ocean meta-virome study reporting that abundances of individual bacteriophages varied significantly across time and space in response to complex deterministic and random processes that influenced infectivity success (Luo, Aylward, Mende, & DeLong, 2017). Considering the communities in halite nodules are largely isolated because of limited inter-nodule exchanges, such dynamic processes could result in the viral activity variance observed in our study, with stochastic factors resulting in unique outcomes in each nodule.

Our study also shed light on the transcriptional functioning of the halite community as a whole. The majority of highly-transcribed pathways in the halite metatranscriptome was related to cell maintenance and basal metabolic activities – transcription, translation, and processes associated with their regulation. The transcriptional landscape indicated rapid carbon turnover by the sole green alga, with its photosynthesis and oxidative phosphorylation pathways being among some of the most consistently highly expressed in the community. In halite nodules, the carbon fixed by *Cyanobacteria* and the alga by way of oxygenic photosynthesis is most likely the only source of primary production (Robinson et al., 2015), as essential genes for ammonia oxidation were not detected in this or previous metagenomes (G. Uritskiy et al., 2019). ATP can also be produced via rhodopsin light-activated proton pumps by heterotrophic *Halobacteria* and *Bacteroidetes* (Engelhard, Chizhov, Siebert, & Engelhard, 2018). We found that opsin production was one of the most upregulated pathways in *Halobacteria*, suggesting that supplementation of their ATP budget by light-driven reactions is essential for these organisms. This is consistent with the upregulation of energy-harvesting rhodopsins in *Halobacteria* during stress (Spudich, 1998) and other studies reporting that opsin-based proton pumps are important supplementary sources of energy for *Halobacteria* in energetically-taxing hypersaline environments (Ernst et al., 2014; Grote & O’Malley, 2011). Finally, it should be noted that while the algae contribute relatively more to the carbon fixation than previously believed, the overall community transcription and metabolic activity rates may still be quite slow, as evidenced by previous estimates of carbon turnover in these microbiomes (Ziolkowski, Wierzchos, Davila, & Slater, 2013).

## Conclusions

Here we report the first characterization of the metatranscriptome of halite microbial communities, providing novel insights into the functioning of this unique ecosystem. We found a surprisingly high variance in the transcriptional landscape of these communities despite a relatively robust functional potential, indicating active transcriptional adaptations to the unique conditions present in each halite nodule. This highly dynamic transcriptional landscape most likely reflects the diversity in rock topology, environmental exposure, and water activity of the nodules. Future studies should be designed to determine the factors regulating this variance on a spatial scale, including a detailed interrogation of halite internal RH and microbiome composition and function. Despite the extreme conditions of the Atacama Desert and the high relative abundance of extremophilic prokaryotes in these endolithic microbiomes, a newly characterized alga was surprisingly found to be the most transcriptionally active member, possibly producing most of the community’s biologically available carbon. Finally, our meta-omic analysis of the community’s metavirome led to the discovery and characterization of several novel and infectively active halophilic viruses and their hosts.

## EXPERIMENTAL PROCEDURES

### Sample collection and processing

Halite nodules were harvested in Salar Grande, a salar in the Northern part of the Atacama Desert (Robinson et al., 2015) in February 2017. All nodules were harvested within a 50m^2^ area as previously described (Robinson et al., 2015) at 9 am and 9 pm, with 6 replicates per time-point, for a total of 12 samples. The colonization zone of each nodules was grounded into a powder, pooling from 3 nodules until sufficient material was collected, and stored in dark in dry conditions until DNA extraction in the lab. At the time of sampling, 4g of powder from each sample was mixed with 4ml of *RNAlater* and stored at 4°C for RNA extraction in the lab. Genomic DNA was extracted as previously described (Crits-Christoph et al., 2016; Robinson et al., 2015) with the DNAeasy PowerSoil DNA extraction kit (QIAGEN). Whole genome DNA sequencing libraries were prepared using the Nextera XT DNA library kit (Illumina) with 1ng of input gDNA. Library amplification was done with dual-index primers for a total of 9 cycles, and the product library was cleaned with XP AMPure Beads (0.6X ratio). Total RNA was extracted from the *RNAlater* samples by first isolating the cells through gradual dissolving of the salt, as previously described (Crits-Christoph et al., 2016; Robinson et al., 2015), cell lysis by mechanical bead beating with the RNAeasy PowerSoil RNA extraction kit (QIAGEN), and extraction from the lysate with a Quick-RNA miniprep kit (Zymo Research); two independent samples were extracted from each replicate. cDNA was generated from 5ng of RNA with the SuperScript III reverse transcriptase (ThermoFisher) using 25 PCR cycles as described previously (Robinson et al., 2015) and the lack of gDNA contamination in the RNA was confirmed by RT-PCR with the 515F/926R 16S rRNA gene primers (Fig. S1). Note that the cDNA was only used for DNA contamination assessment, but not library construction. RNAseq libraries were prepared with the *SMARTer* Stranded RNA-seq kit (*TaKaRa*) using 25ng of RNA input and 12 cycles for library amplification. All other steps followed the manufacturer’s recommendations. 24 paired RNA libraries corresponding for the 12 metagenomic samples were pooled *in-silico* (files were concatenated) into 12 replicates to exactly match the sequenced material in the metagenomic samples. The final barcoded libraries were quantified with Qubit dsDNA HS kit, inspected on a dsDNA HS Bioanalyzer, pooled to equal molarity, and sequenced with paired 150bp reads on the HiSeq 2000 platform at the Johns Hopkins Genetic Resources Core Facility (GRCF).

### Processing shotgun metagenomic and metatranscriptomic sequence data

The de-multiplexed shotgun reads were processed with the metaWRAP v1.1 pipeline (G. V. Uritskiy, DiRuggiero, & Taylor, 2018) with recommended databases on a UNIX cluster with 112 cores and 2048GB of RAM available. Read trimming and human contamination removal was done by the metaWRAP Read_qc module (default parameters) on each sample. The metatranscriptomic reads were digitally ribo-reduced with SortMeRNA v2.1b (Kopylova, Noe, & Touzet, 2012) by aligning the reads to SILVA v138 ribosomal sequences. The reads from all metagenome replicates were co-assembled with the metaWRAP Assembly module (--use-metaspades option) (Nurk, Meleshko, Korobeynikov, & Pevzner, 2017). Each replicate was also assembled individually for algae sequence extraction (described below). For metagenome-assembled genome (MAG) recovery, the co-assembly was binned with the metaWRAP Binning module (--maxbin2 --concoct --metabat2 options), and the resulting bins were then consolidated into a final bin set with metaWRAP’s Bin_refinement module (-c 70 -x 5 options). The total abundances (with metagenomic reads) and total expression (with metatranscriptomic reads) of MAGs and contigs were then quantified in each replicate by Salmon (Patro, Duggal, Love, Irizarry, & Kingsford, 2017) with the Quant_bins module (default parameters). All scripts and intermediate data used for this analysis are publicly available at https://github.com/ursky/metatranscriptome_paper.

### Functional and taxonomic annotation

Gene prediction and functional annotation of the co-assembly was done with the JGI Integrated Microbial Genomes & Microbiomes (IMG) (I. A. Chen et al., 2017) annotation service. Taxonomy of each contig was estimated by computing an average of the taxonomic annotation of its genes, as returned by the IMG service. The taxonomic depth was reduced until >50% of gene taxonomies agreed. Gene relative abundances in the metagenomes were taken as the average DNA read depth of the contigs carrying those genes and expressed as copies per million reads (CPM). Gene expression was estimated with Salmon (Patro et al., 2017) and expressed as transcripts per million reads (TPM). KEGG KO identifiers were linked to their respective functions using the KEGG BRITE pathway classification (Kanehisa, Sato, Kawashima, Furumichi, & Tanabe, 2016). For the functional annotation of pathways in specific taxa or MAGs, only pathway with minimum of 5 unique enzymes, constituting at least 20% of all possible enzymes in that pathway, were used. KEGG pathway total abundance (from metagenomes) and total expression (from metatranscriptomes) were calculated as the sum of the TPMs or CPMs of genes classified to be part of the pathway. When comparing the total pathway abundance in the metagenomes to their total expression in the metatranscriptomes, the abundance and expression values were standardized to the sum of the CPMs or TPMs of the pathways in each sample, respectively. All scripts and intermediate data used for this analysis are publicly available at https://github.com/ursky/metatranscriptome_paper.

### Metatranscriptomic Differential Expression Analysis

Gene expression values from Salmon (Patro et al., 2017), expressed in transcripts per million reads (TPM), were further standardized in each sample to the average abundance (CPM) of the organism carrying each gene, evaluated at the contig or MAG level. In an alternate test, the expression data was standardized to the total average expression of each contig or MAG. The standardized expression values were loaded into DESeq2 (Love, Huber, & Anders, 2014) to identify differentially expressed genes between the two test groups – samples collected at 9 am and 9 pm. This analysis was repeated for the whole community (using only contigs longer than 5kb for robust standardization) and individual MAGs, particularly the *Halothece* genome, the *Dolichomastix* cellular and organelle genomes. For genes that were found to be putatively enriched in the 9 am or 9 pm time points (DESeq adjusted p-value<0.01), pathway enrichment analysis was performed by comparing the pathway functions of the “morning” and “evening” genes to that of all the genes in the community with a sub-sampling simulation (n=10,000).

### *Dolichomastix* alga genome extraction

The main genome of the alga was extracted from the co-assembly with metaBAT2, which yielded a 10.3Mbp main chromosomal genome (N50=6.0kb). The binning accuracy was manually assessed by interrogating the phylogeny of the genes carried on the binned contigs, as determined with the IMG functional annotation, above. Due to extremely high coverage of the chloroplast and mitochondrion (∼140X and ∼70X, respectively), their sequences could not be directly extracted from the co-assembly. A few initial short contigs were extracted from the co-assembly by aligning with BLAST v2.6.0 to previously identified contigs of the halite alga chloroplast and mitochondrion (Crits-Christoph et al., 2016). These sequences were then manually curated based on taxonomy and coverage and aligned to the individual replicate assemblies to identify longer and more accurate contigs. The completion of the genome was estimated with BUSCO v3.1.0, using the chlorophyta_odb10 lineage database (Seppey et al., 2019). Using metaWRAP’s reassemble_bins module, all chromosomal and mitochondrial contigs from the individual assemblies were used to pull out reads from all the samples, and the corresponding reads were reassembled into final sequences. The chloroplast sequence was 95Kbp (N50=64.5Kbp) and the mitochondrion sequence was 46Kbp (N50=16.7Kbp). The corresponding contigs in the main co-assemblies were replaced with these sequences and re-annotated with IMG.

### *Dolichomastix* genome analysis

To investigate predicted proteome adaptations, the isoelelectric point (p*I*) of proteins predicted from genes of several algae were compared. The chromosomal genomes of the extracted *Dolichomastix* genome and 3 other algae – *Dunaliella salina*, *Ostreococcus tauri*, and *Micromonas pusilla* – were annotated with GeneMarkS (Besemer, Lomsadze, & Borodovsky, 2001) and the “intronless eukaryotic” setting, and the amino acid sequences of their genes analyzed with ProPAS v1.1 (Wu & Zhu, 2012). To compare pI values of homologous proteins, the translated genomes of *Dolichomastix* and *M. pusilla* were aligned with BLAST v2.6.0 to identify paired values. A minimum percent identity of 40%, alignment length 50nt, coverage of 40%, and a maximum e-value of 0.01 (full command: blastp -query dunaliella_cellular_proteins.faa -subject Dolichomastix.faa -outfmt “6 qseqid sseqid pident length mismatch gapopen qstart qend sstart send evalue bitscore qcovhsp” -max_target_seqs 1 > dunaliella_cellular_proteins.blast) were required for a pair of proteins to be considered homologous, and multiple hits were de-replicated to only consider the best hit (minimum e-value). To identify genes homologous to important genes in the *D. salina* genome, the amino acid sequences of the genes were similarly aligned to the *Dolichomastix* genes to identify homologues.

### Viral contig extraction and annotation

Viral sequences were pulled out from the co-assembly with *de-novo* non-targeted viral sequence discovery (Paez-Espino, Pavlopoulos, Ivanova, & Kyrpides, 2017). The abundance (CPM) and total expression (TPM) of the viral contigs was compared between replicates, and Pearson coefficient was computed for the correlation of abundance and expression of viruses across the 12 replicates. For each genus-level group from VContact2 (Bin Jang et al., 2019), genomes and protein sequences were extracted. To determine the best genes for phylogenetic tree construction, All-vs-all comparison of protein sequences was conducted using EFI-EST (Gerlt et al., 2015), and the most conserved gene present as a single paralog was used for tree construction. Given the fragmentary nature of some contigs from this study, not every genome could be compared with a single marker gene. Alignment of marker protein sequences was conducted with PROMALS3D (Pei, Kim, & Grishin, 2008). IQ-Tree was used to make phylogenetic trees of alignments, with best substitution model automatically determined, and 1000 ultrafast bootstraps used for each tree. Genome map comparisons were done in EasyFig (Sullivan, Petty, & Beatson, 2011) using TBLASTX alignment at E Value < 0.01. For host determination of sequences from this study, CRISPR spacers extracted from the metagenomic assembly with MinCED were matched to viral contigs using BLASTN (Altuschul, Gish, Miller, & Lipman, 1990).

## ACKNOWLEDGMENTS

**General:** We thank Samantha Getsin and Sean Ravel for help in sample processing.

## Author contributions

GU, JT, and JD conceptualized and designed the study; GU, DRG, and JD collected in-field samples; GU and AM processed and sequenced samples; GU analyzed the data and wrote the manuscript; JT and JD edited the manuscript.

## Competing interests

The authors declare no competing interests.

### Data availability

- Raw metagenomic and metatranscriptomic sequences are publicly available through NCBI PRJNA560058.
- Metagenomic assembly and its functional annotation are publicly available through the GOLD service at JGI (Ga0371442, Taxon ID 3300033522).
- All intermediate data, MAGs, and analysis scripts used for this analysis are publically available at https://github.com/ursky/metatranscriptome_paper.
- Supplementary material available online.

## SUPPLEMENTARY MATERIAL

**Fig. S1:**
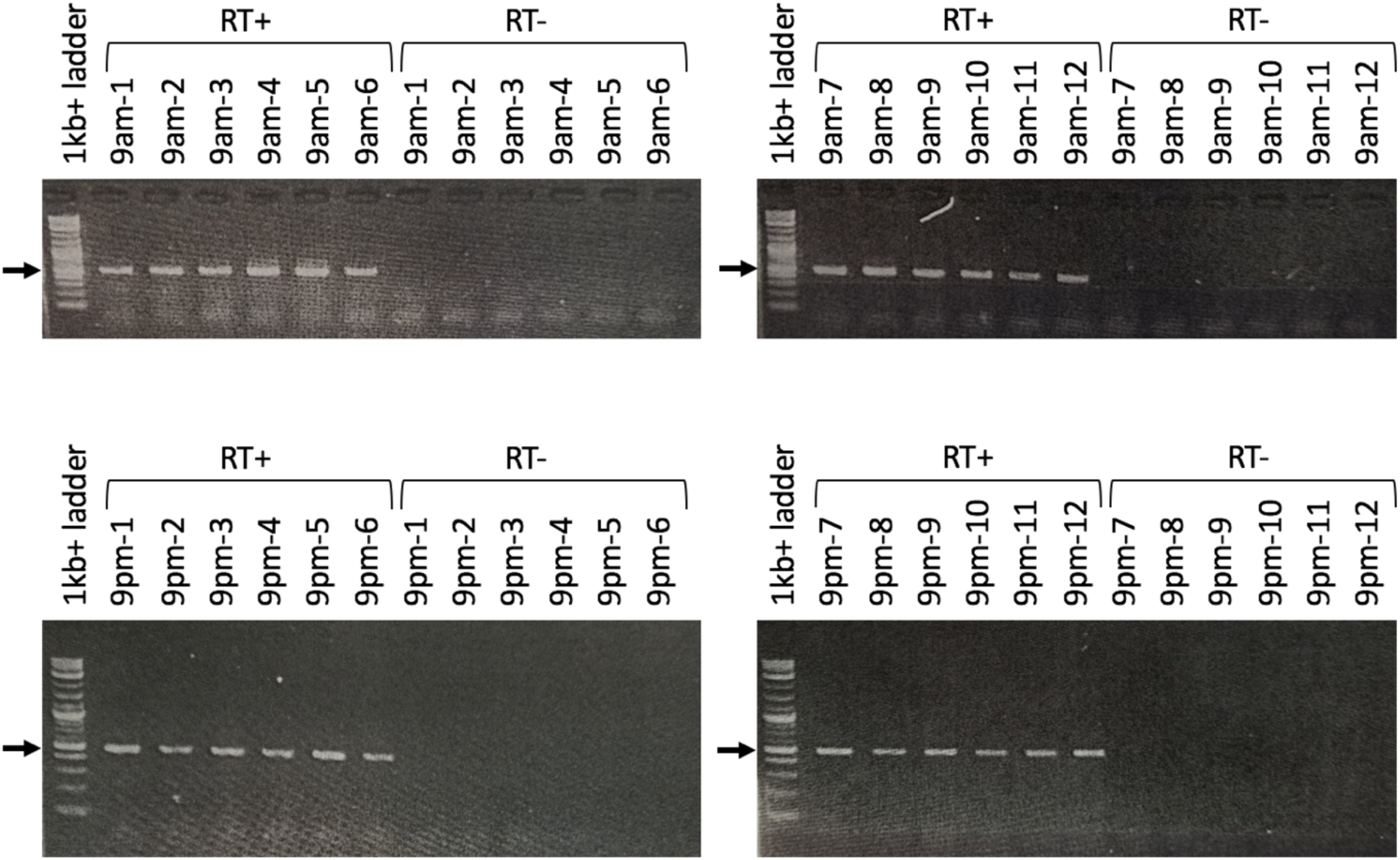
RT-PCR validation of the lack of DNA contamination in the extracted RNA. Each RNA sample was processed using the SuperScript III cDNA synthesis kit with (RT+, positive control) or without (RT-, negative control) the reverse transcriptase enzyme. The 16S rRNA gene was amplified from the resulting cDNA. Pairs of neighboring samples (1 and 2, 3 and 4, 5 and 6, etc.) were pooled in-silico to produce the DNA metagenomic libraries. Arrows indicate the expected locations of amplicon bands (512bp).

**Fig. S2:**
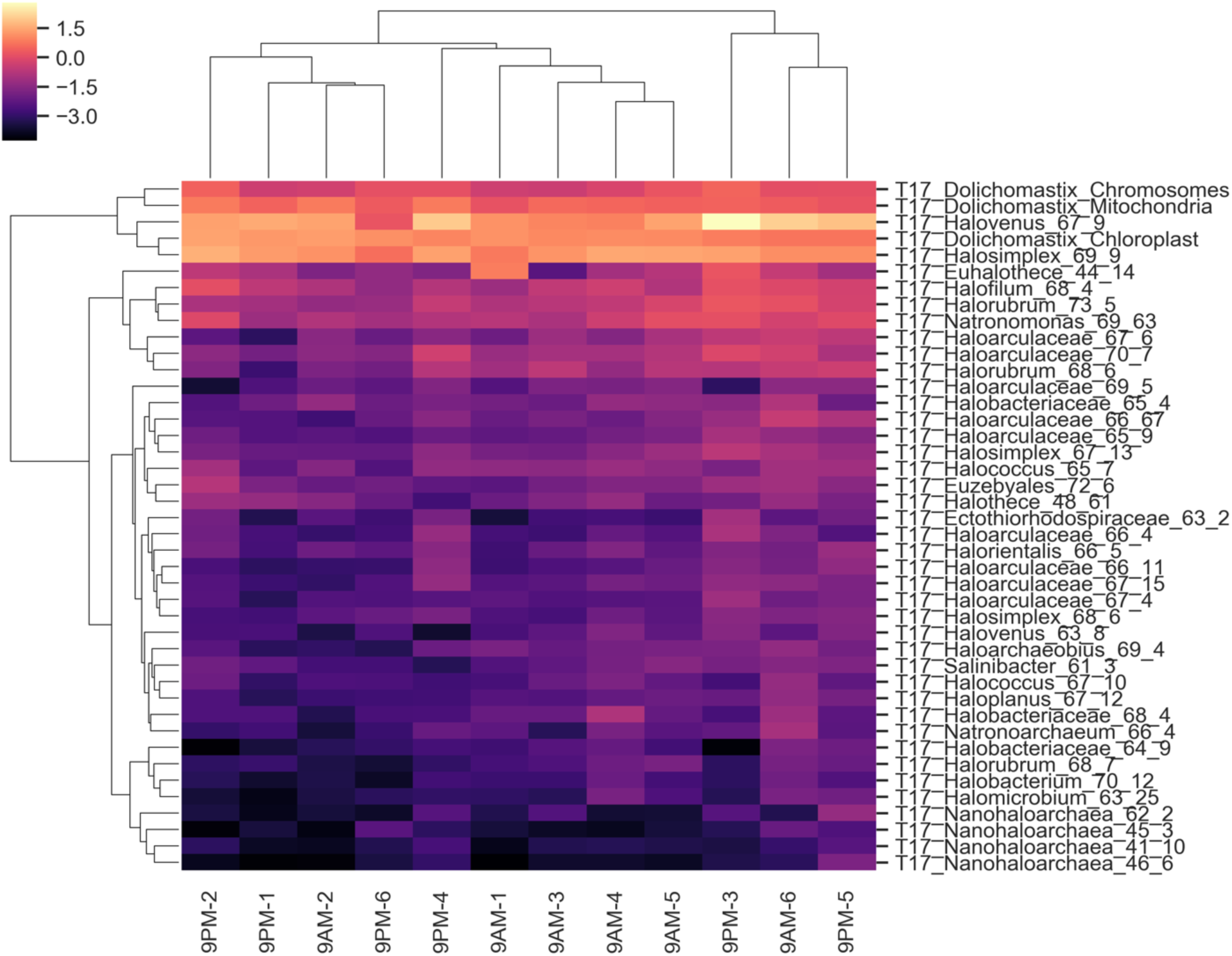
Log values of ratios of the average transcription (TPM) over the average abundance (CPM) for high-quality MAGs (>70% completion, <5% contamination) across the 12 replicate samples.

**Fig. S3:**
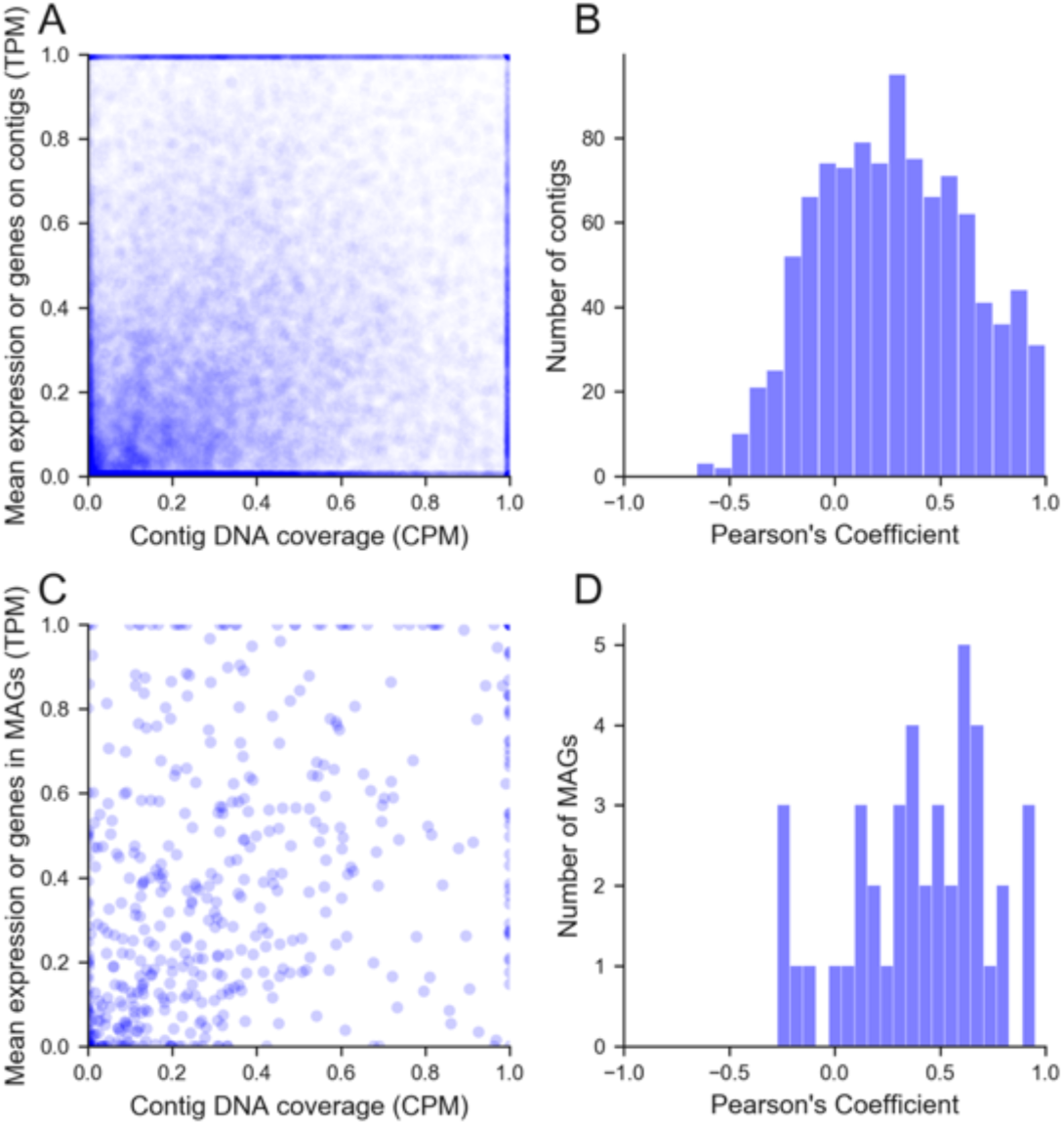
Relationship between scaled contig (A) and MAG (C) mean expression and scaled abundance in all replicates. The scaled values were calculated as follows: TPM_scaled_=(TPM-TPM_min_)/(TPM_max_-TPM_min_), CPM_scaled_=(CPM-CPM_min_)/(CPM_max_-CPM_min_). Distributions of Pearson correlation coefficients in contigs (B) and MAGs (D) of the non-scaled TPM and CPM values across the 12 replicates.

**Fig. S4:**
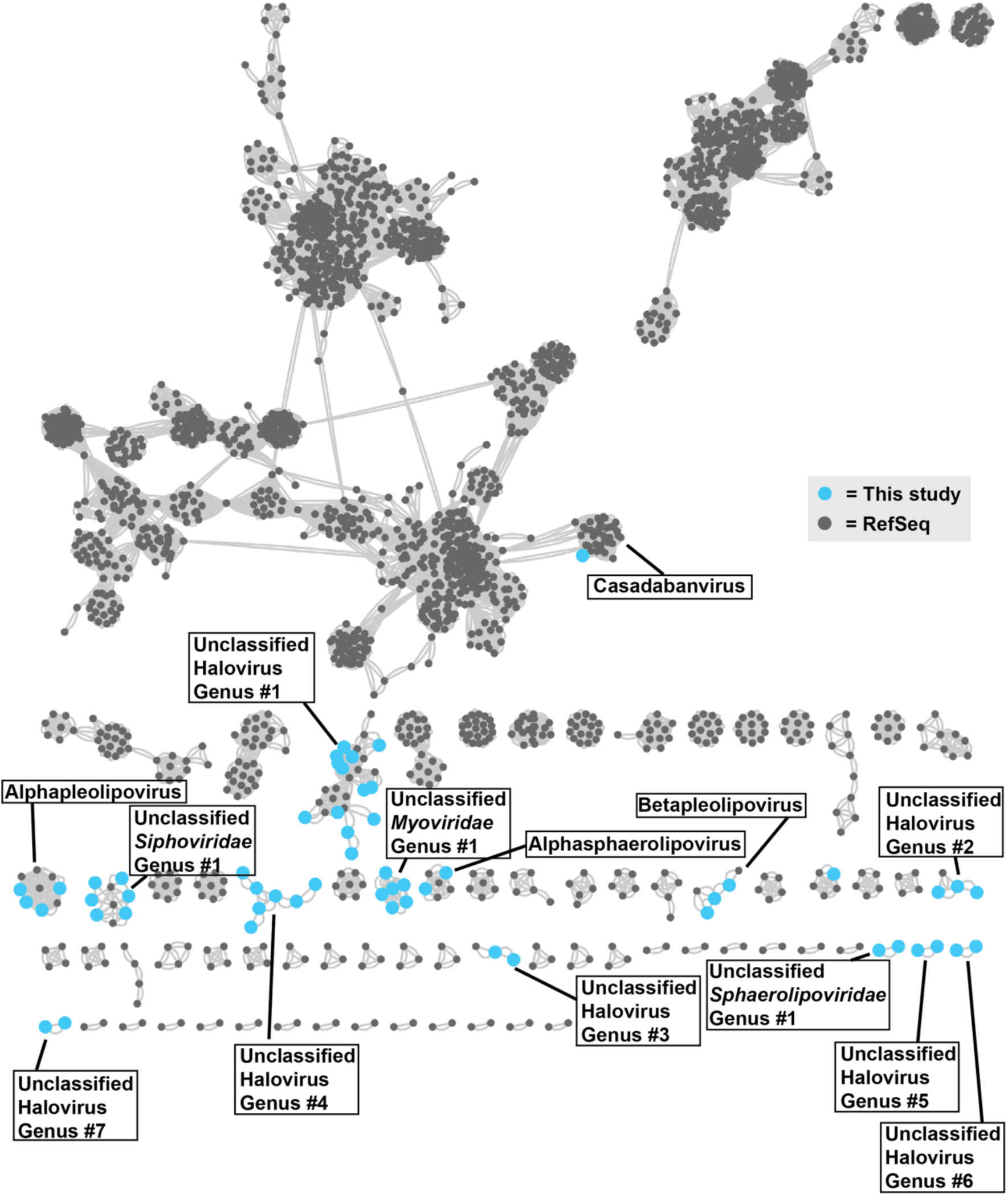
Gene similarity network of RefSeq viruses (grey) and with viral contigs identified in this study (blue). Genus-level taxonomy was determined using VContact2 and inter-genome similarity was estimated with protein alignments with DIAMOND. Any genomes with no sufficient similarity to other genomes were not displayed.

**Fig S5:**
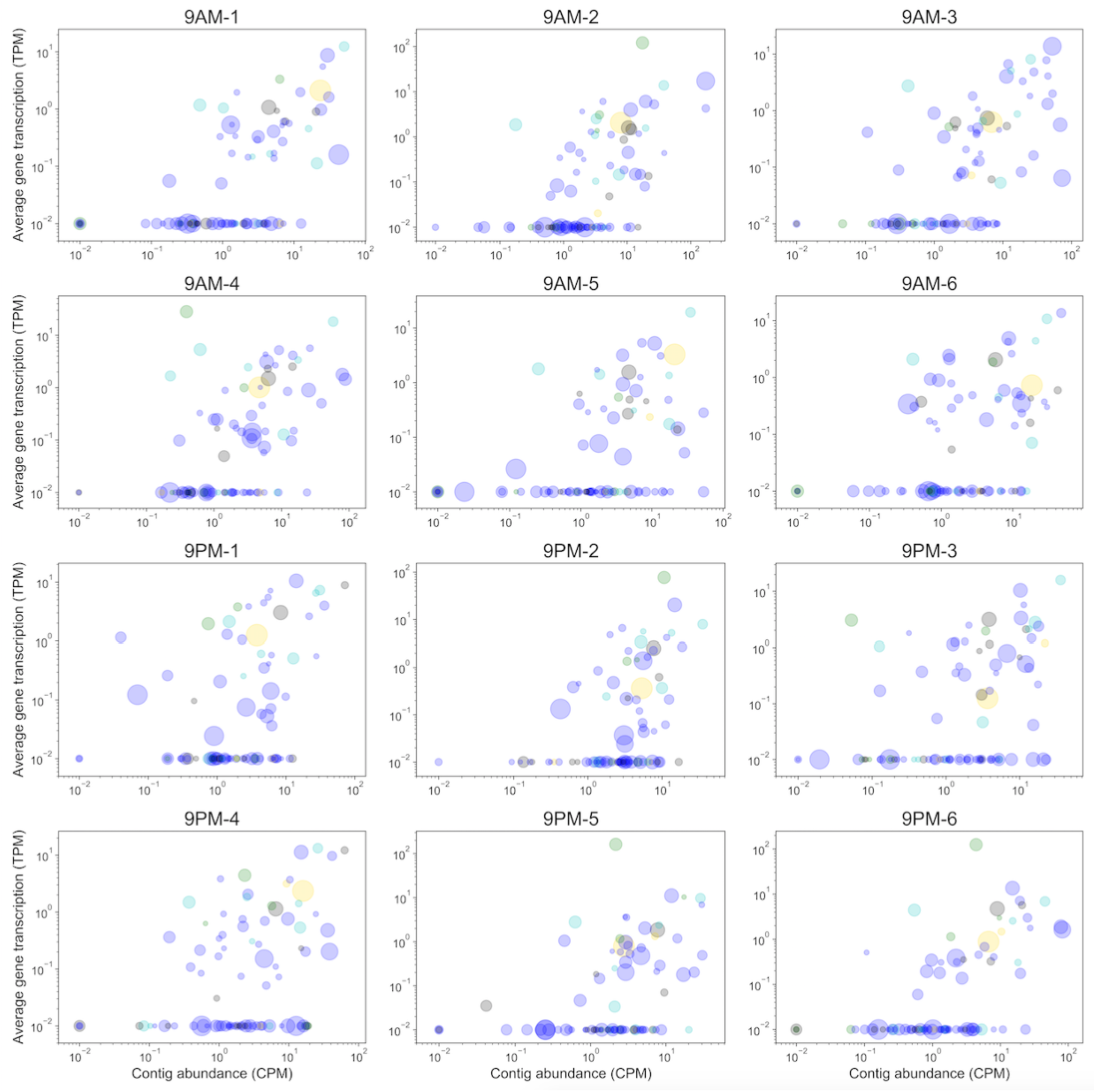
Viral contig total transcriptional activity (TPM) and abundance (CPM) across the 12 replicates. Size of circles represents the relative length of viral contigs and color denotes the taxonomy of predicted host (blue: Euryarchaeota, cyan: Proteobacteria, green: Cyanobacteria, gold: Bacteroidetes, gray: unknown).

**Fig. S6:**
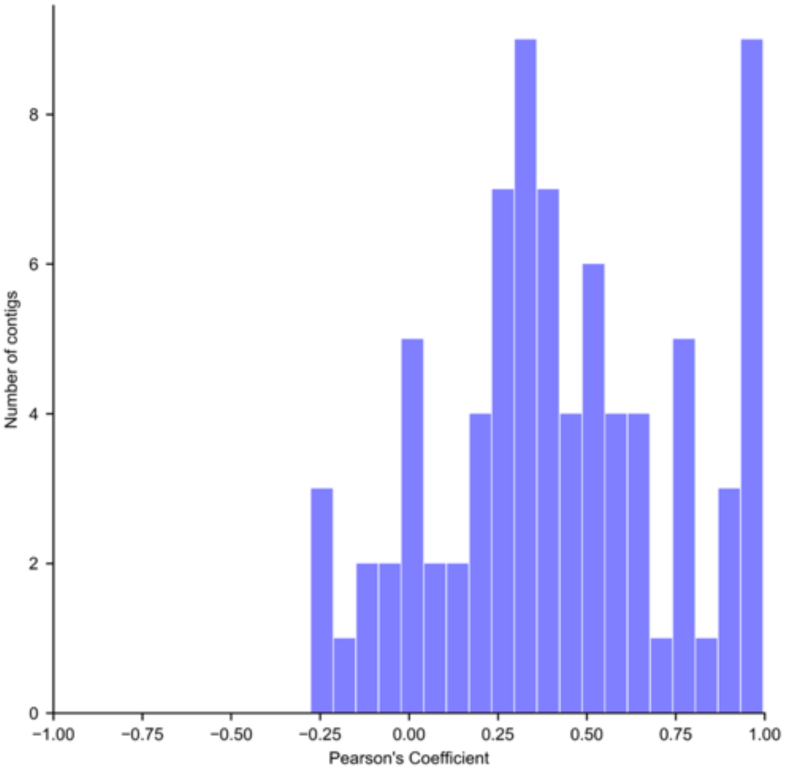
Distribution of Pearson correlation between total transcriptional activity (TPM) and total abundance (CPM) of viral contigs across the 12 replicates.

**Fig S7:**
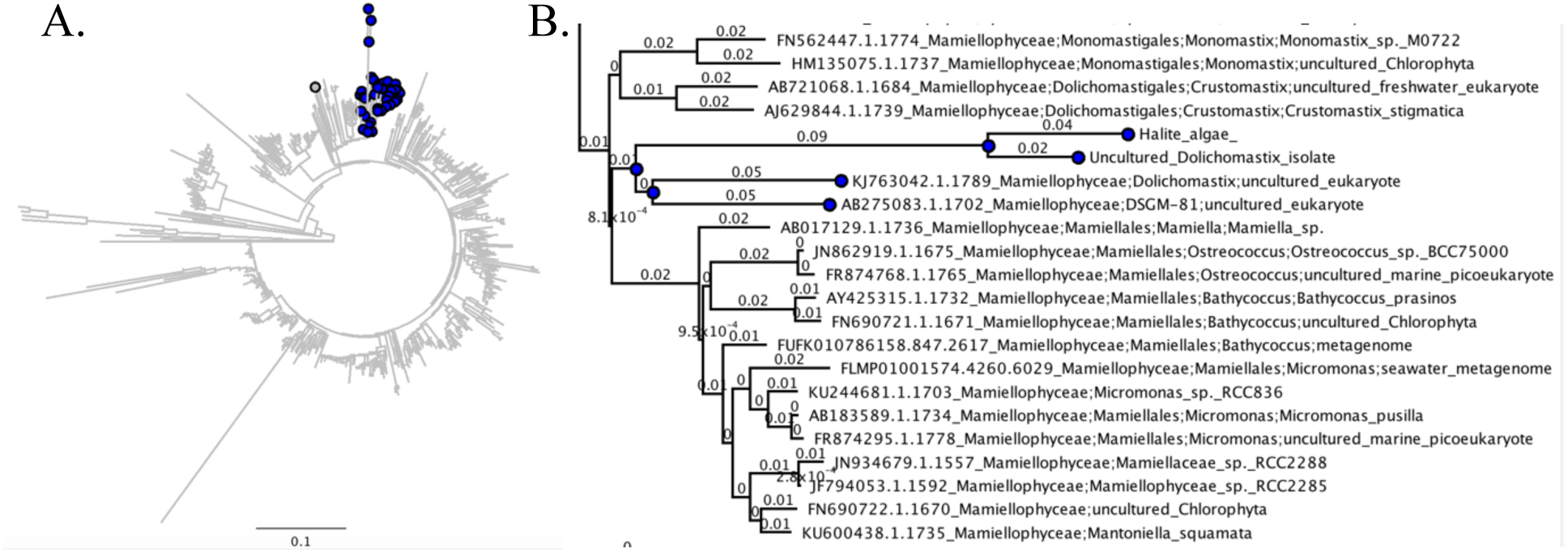
(A) Unrooted phylogenetic tree of a random subset of 500 Chlorophyta (green algae) species from the SILVA database, including all available sequences from the Mamiellophyceae class (highlighted in blue), constructed from a multiple alignment of 18S rRNA sequences. (B) Zoomed-in view of the Mamiellophyceae class phylogeny, showing the clustering of 18S rRNA gene sequences from the halite algae with members of the Dolichomastix genus (highlighted in blue; the closest sequence from Lake Tyrrel isolate under GenBank accession number KC486366.1).

**Fig. S8:**
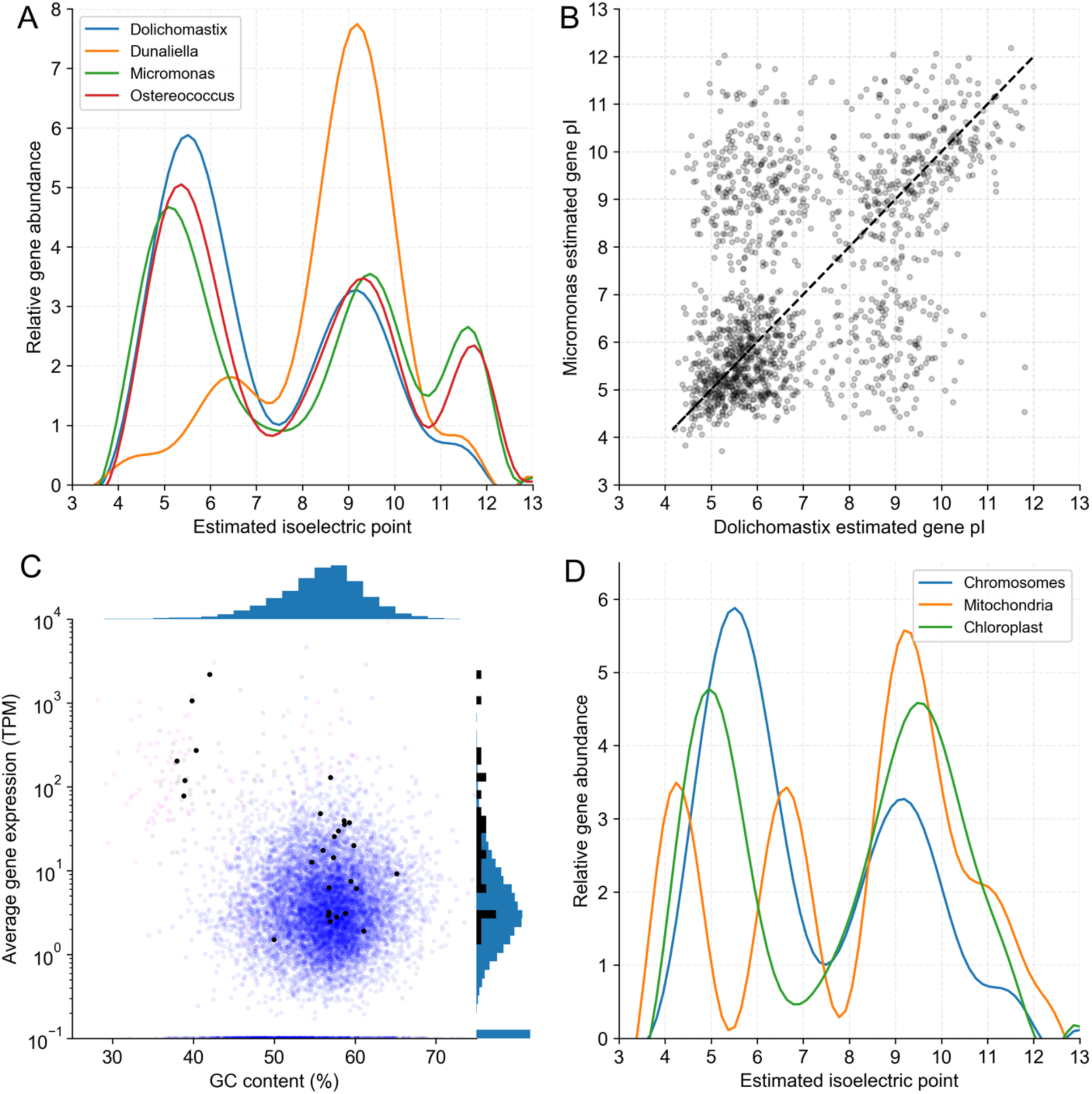
(A) Distribution of isoelectric points (pI) of predicted proteins in various algal genomes, including the novel Dolichomastix. (B) Predicted pIs of homologous proteins identified in both Dolichomastix and M. pusila genomes, showing biases in pI preferences between the two genomes. (C) GC content and average expression of Dolichomastix genes, with genes encoding homologs of proteins important for high-salt tolerance in D. salina highlighted in black. (D) Distribution of isoelectric points (pI) of predicted genes in Dolichomastix chromosomes, mitochondria, and chloroplast.

**Fig. S9:**
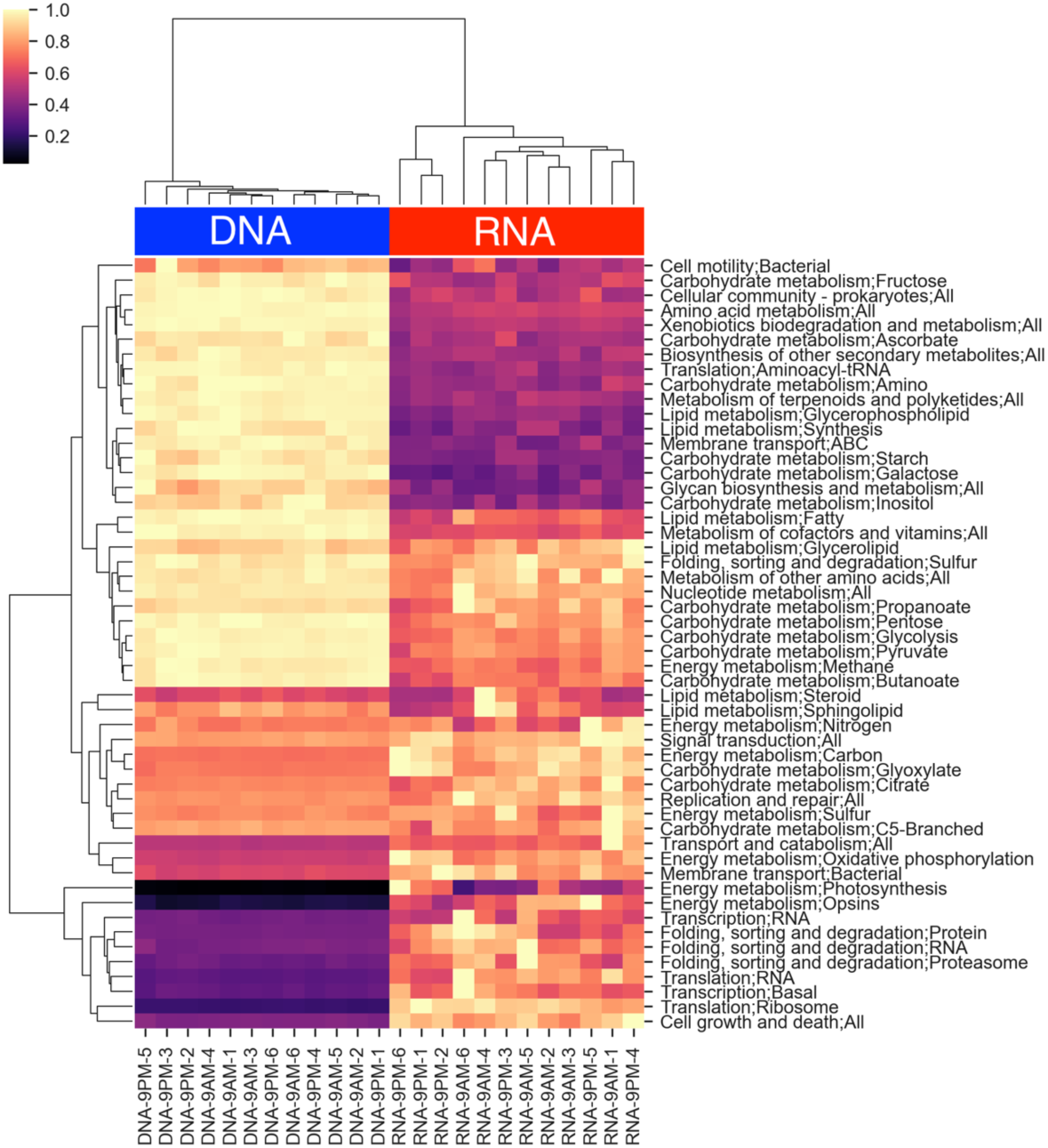
Relative prevalence of major KEGG functional pathways in the metagenomic and metatranscriptomic samples. The pathway abundance (DNA) and expression (RNA) value were the combined total CPM and TPM, respectively, of the genes that constituted that pathway. All samples were standardized to an equal total coverage of all pathways, and then scaled to the maximum value for each row (pathway).

**Fig. S10:**
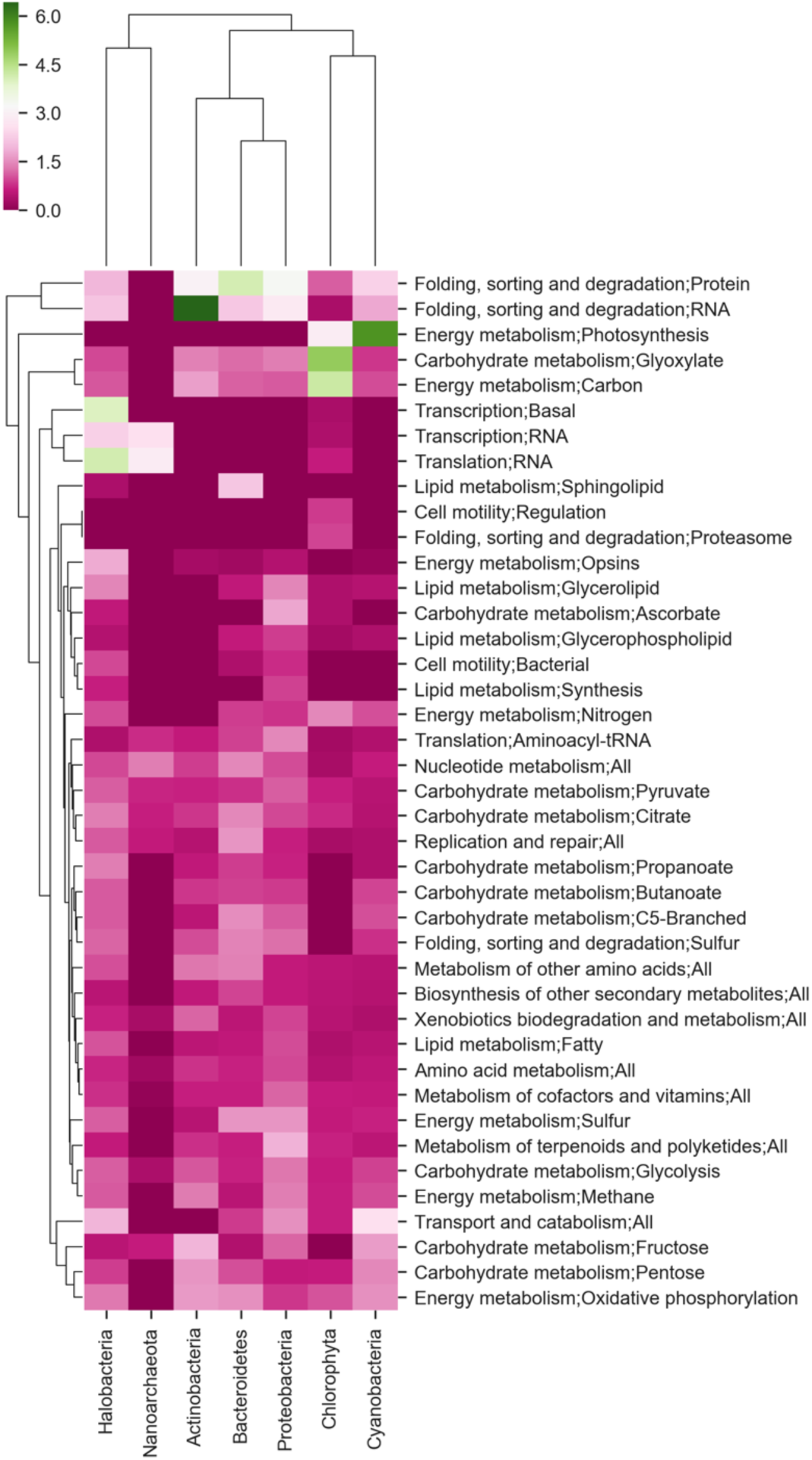
Mean transcriptional activity ratio of major highly-expressed KEGG pathways in the major taxa found in the halite communities. The relative activity is the ratio of total pathway expression (TPM) to the pathway’s abundance (CPM) in each taxon.

**Fig S11.**
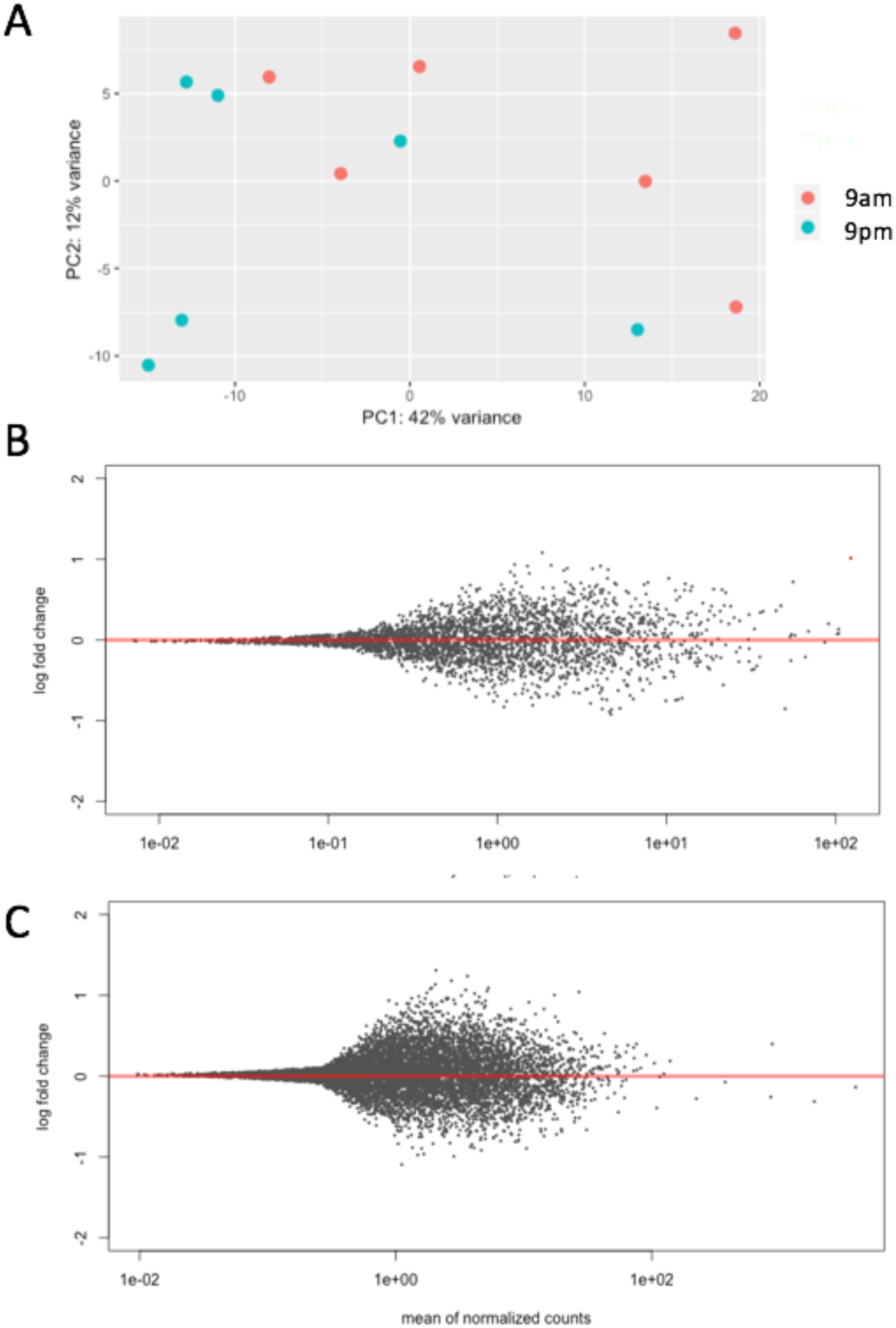
DESeq differential expression analysis of the halite community between the 9 am and 9 pm time points. (A) Clustering between the time points shown with a PCA of the expression profile of the entire community. (B) Lack of significantly differentially expressed genes (q-value<0.01) in the Halothece and (C) Dolichomastix MAGs. Gene expression was expressed as transcripts per million (TPM) and further standardized to the total transcription of the contig (A) or MAG (B,C) that they belong to in order to account for inter-replicate variation in abundance and activity.

**Fig. S12:**
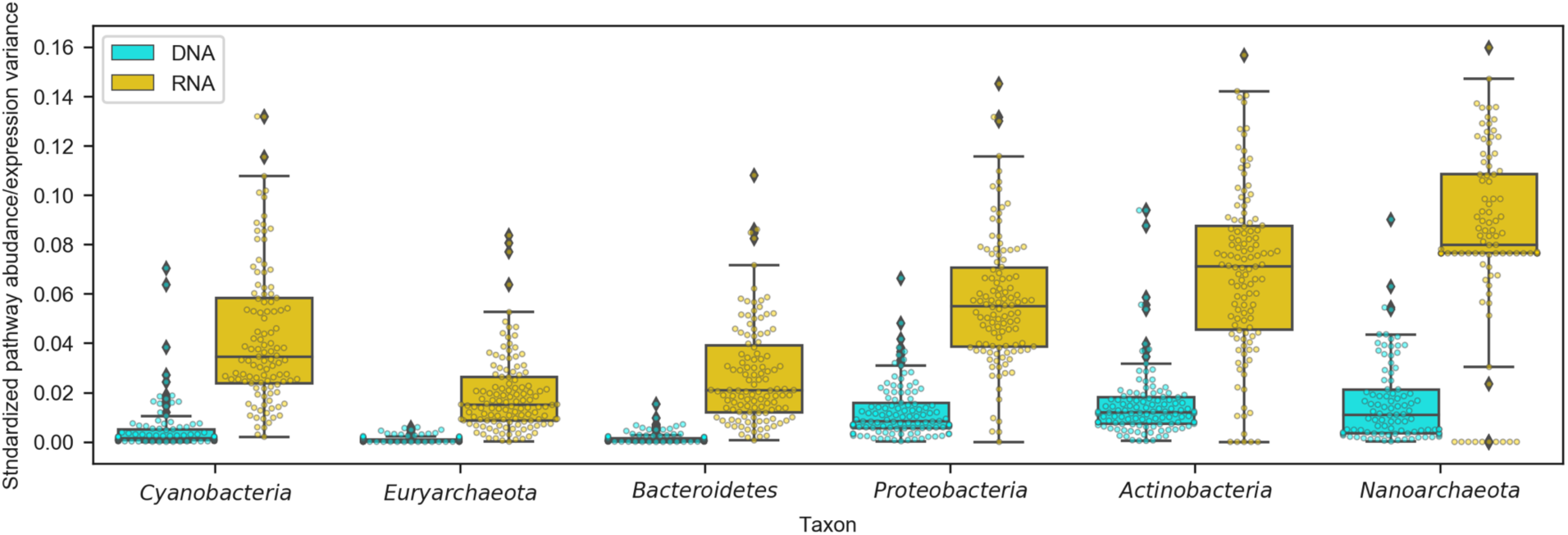
Variance in pathway abundance in metagenome replicates (cyan) and variance in pathway expression in metatranscriptome replicates (gold) shown for major phylogenetic groups of the halite community.

